# Inhgrams for engrams: co-active inhibitory-inhibitory plasticity shapes inhibitory assemblies that stabilize and recall embedded engrams through disinhibition

**DOI:** 10.64898/2026.09.15.751748

**Authors:** Maciej Kania, Basile Confavreux, Tim P. Vogels

**Affiliations:** Institute of Science and Technology Austria, Klosterneuburg, Austria; Gatsby Computational Neuroscience Unit, UCL, London, UK

## Abstract

Inhibitory neurons have been widely understood to play a supporting role in neural function and memory formation, but recent advances highlight that memories are encoded by both excitatory and inhibitory neurons (EI assemblies), carving out a bigger role for inhibition beyond mere stabilization. However, the computations enabled by such EI assemblies are still unclear, and the synaptic plasticity rules that could sustain and retrieve memories are unknown. Here, we construct a computational model for EI assembly recall and investigate the computational benefits of such mixed engrams over classical, excitatory-only engrams. Towards this end, we consider large recurrent spiking networks with symmetrical Hebbian synaptic plasticity at both inhibitory-to-inhibitory (I-to-I) and inhibitoryto-excitatory (I-to-E) synapses. The conjunction of these rules can robustly stabilize embedded EI assemblies in the asynchronous irregular regime. Assemblies can then be reactivated by two distinct mechanisms: direct stimulation of the engram or disinhibition through the inhgram. Both mechanisms of recall lead to reliable pattern completion and separation. Crucially, we show that networks that lack I-to-I plasticity show weak recall and cannot discriminate between overlapping engrams via disinhibitory activation. This suggests that inhibitory plasticity can facilitate recall of overlapping engrams, with inhibitory neurons controlling multiple excitatory engrams. Furthermore, we show that inhgrams can emerge from pre-existing E-I-E loops in the network’s random connectivity. Our work proposes that inhibitory plasticity is a plausible mechanism for high-quality disinhibitory recall, allowing inhibitory neurons to selectively control multiple excitatory engrams through synapse-specific plasticity rules.

**AUTHOR SUMMARY:** The brain is thought to store memories in the form of engrams, groups of neurons that reactivate together during memory recall. Until recently, research focused almost exclusively on excitatory neurons as the building blocks of these memory traces, while inhibitory neurons were assumed to play a supporting role, responsible only for keeping network activity stable. However, recent experimental work has challenged this notion by showing that inhibitory neurons appear to form their own memory-related assemblies (inhgrams). In fact, manipulating inhibition in humans and animals can unmask or erase memories. This raises a fundamental question: what synaptic mechanisms could support such excitatory-inhibitory memory assemblies, and how would the brain use them to retrieve memories? Here, we used simulations of large spiking networks of excitatory and inhibitory neurons to show that when inhibitory synapses undergo plasticity, memories can be selectively retrieved by silencing inhibitory neurons, even when multiple memories partially overlap. Moreover, inhibitory plasticity amplifies pre-existing connectivity motifs, making the inhibitory component of a memory predictable before it is even formed. Our results suggest that inhibitory neurons are active organizers of memory retrieval, not just stabilizers, and play an important role in learning how to discriminate between similar memories.

## I. INTRODUCTION

Memories are thought to be encoded and stored in the form of *engrams*, interconnected (excitatory) cell assemblies representing a memory trace and reactivated during memory recall^1^. However, in light of work on the role of precise excitatory-inhibitory (EI) balance^2–6^ and inhibitory plasticity in neural networks^7–10^, the concept of *inhgrams*, i.e., groups of inhibitory neurons balancing the activity of strongly interconnected engrams, has been proposed^11,12^. In line with this idea, the manipulation of local GABA levels has been demonstrated to unmask memories in human subjects^13^. Further, the activation of disinhibitory circuits in rodents has been shown to enable recall of fear memories^14,15^; and perturbed inhibition can reopen the critical window of song learning in zebra finches^16^.

Growing evidence suggests that excitatory and inhibitory neurons (or engrams and inhgrams) may come in strongly coupled subnetworks, referred to as EI assemblies^17,18^. Such assemblies provide computational benefits beyond network stabilization, including stimulus-specific computations^17^, structured and balanced neural representations^19^, and associative-memory processing^18^. However, little is known about how EI assemblies enable memory recall, or what synaptic mechanisms could support the formation, stabilization, and retrieval of their inhibitory component. Inhibitory synaptic plasticity emerges as a natural candidate^9,20,21^, acting to strengthen or weaken synapses onto excitatory neurons and thus helping to form inhgrams. Additional plastic control of inhibitory-to-inhibitory (I-to-I) synapses could help to link individual inhgram neurons, allowing them to form assemblies of their own (inhgrams) through shared disinhibition.

Whereas prior work treats inhibition as balancing or gating fixed engrams, here we let both inhibitory-to-excitatory and inhibitory-to-inhibitory synapses learn, and ask what this co-active (I-to-All) plasticity buys for memory recall. Towards this end, we manually embedded EI assemblies in spiking network models by strengthening the appropriate E-to-E and E-to-I synapses between engrams and inhgrams, respectively. We could demonstrate two successful mechanisms of engram reactivation through direct stimulation, or through indirect disinhibitory action.

Further, networks with plastic I-to-I connections proved more sensitive to pattern-complete partial cues, and showed better recall performance compared with I-to-E-only networks. When we tested for pattern separation, only EI assemblies embedded in the I-to-All network could reliably discriminate between overlapping engrams. Finally, we show that I-to-All plasticity took advantage of pre-existing E–I–E loops in the random connectivity of the network, making it possible to predict which neurons became an inhgram after engram neurons were selected.

## II. RESULTS

Experimental efforts to pinpoint the role of inhibitory mechanisms in memory formation and maintenance remain difficult. To investigate the potential benefits of combined EI assemblies in neural networks, we turned to artificial neural networks, where we aimed to show that co-active inhibitory synaptic plasticity is a plausible and well-suited candidate for this function.

### Stability of neural networks with inhibitory-to-all synaptic plasticity

We started our investigation by exploring network stability with two simultaneously active inhibitory plasticity rules in a recurrent spiking neural network of 10,000 excitatory and inhibitory leaky integrate-and-fire (LIF) neurons^22^ (Fig. 1A). Previous studies established that I-to-E synapses could follow a symmetrical, activity-dependent plasticity rule enforcing precise and detailed balance^22,23^. For simplicity, we used the same rule for the I-to-I connections (Methods).

**Fig 1.**
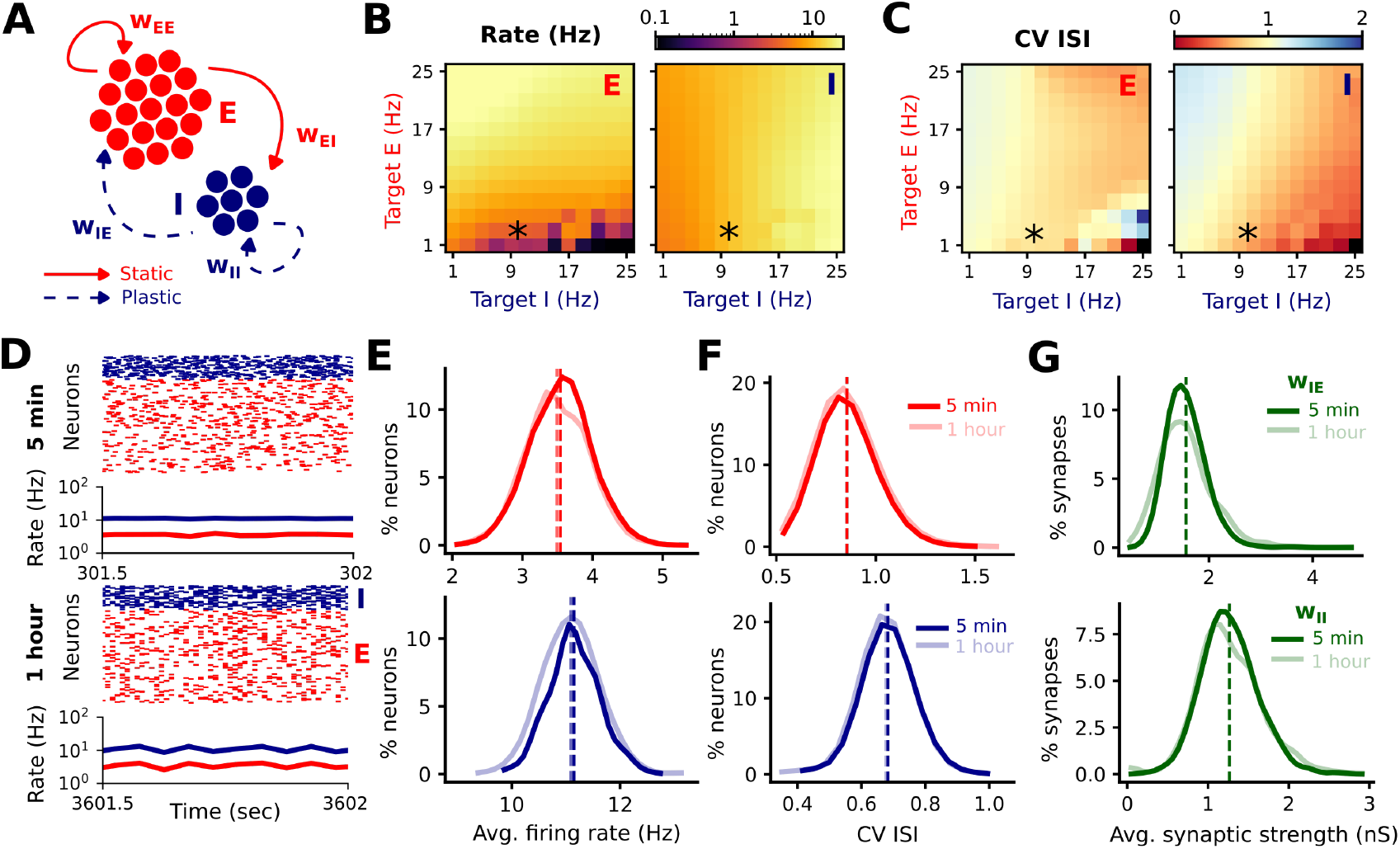
Co-active inhibitory plasticity drives long-term stable network activity. **A:** Schematic of a recurrent spiking network (RSN) with static excitatory and plastic inhibitory connections. **B:** Firing rates of excitatory (left) and inhibitory (right) populations as a function of target E and I firing rates after 5 min of simulation; asterisks indicate the network pursued throughout the study. **C:** Coefficient of variation of the interspike interval (CV ISI) for excitatory (left) and inhibitory (right) populations across the same target rate grid; asterisks as in B. **D:** Raster plots of 400 excitatory (red) and 100 inhibitory (blue) neurons and corresponding population firing rates over 0.5 seconds, shown at 5 minutes (top) and 1 hour (bottom) of simulation. **E:** Distributions of average firing rates across neurons at 5 minutes (dark) and 1 hour (light) for E (top, red) and I (bottom, blue) populations; dashed lines indicate population means. **F:** Distributions of CV ISI at 5 minutes and 1 hour for E (top) and I (bottom) populations. **G:** Distributions of average synaptic strengths (nS) for *w*_*IE*_ (top) and *w*_*II*_ (bottom) connections at 5 minutes and 1 hour; dashed lines indicate means.

To define the regime in which the combination of these rules yields stable network activity, we performed a parameter sweep over the target firing rates of the excitatory (*ρ*_0*E*_) and inhibitory (*ρ*_0*I*_) populations (Methods). We computed average firing rate (Fig. S1A, Fig. 1B), mean squared error (MSE) between target and measured firing rate (Fig. S1B), average interspike intervals (ISIs) (Fig. S1C), coefficients of variation (CV ISI) (Fig. S1D, Fig. 1C), and the average synaptic strengths of both I-to-E and I-to-I connections over 10 s after an initial burn-in period of 5 min (Fig. S1E). We found a range of stable solutions for asynchronous, irregular spiking activity. These solutions occurred across all tested values of *ρ*_0*E*_, however, they were limited across *ρ*_0*I*_ values. We found stable regimes for inhibitory target firing rates (*ρ*_0*I*_) ranging from 1 to 49 Hz.

We chose an example network with low firing rates (*ρ*_0*E*_ = 3 Hz and *ρ*_0*I*_ = 10 Hz) and went on to investigate its long-term stability. We computed the same metrics and their distributions for two time points—5 min and 1 hour (Fig. 1D-G). We show that combined inhibitory plasticity produces cortical-like, asynchronous, and irregular spiking activity that remains stable for at least 1 hour (simulations were terminated after 1h simulated time). Population firing rates remained slightly above the target values (Fig. 1E), but this did not affect stability, as firing rates were unchanged between 5 min and 1 hour of simulation.

### Modelling excitatory-inhibitory assembly recall

Motivated to study memory recall mediated by disinhibitory circuits^14,15^, we embedded an excitatory-inhibitory (EI) assembly in the network (Fig. 2AB) and waited for 5 min simulated time for the network to reach steady activity (Fig. 2C). Next, we manually strengthened the E-to-E and E-to-I connections between 784 excitatory and 196 inhibitory neurons (9.8% of each population^22^). Strengthening excitatory connections resulted in elevated assembly activity. The rest of the population activity remained near-unaffected (Fig. 2D). After 5 min the network returned to global stability (Fig. 2E), such that the activity of the EI assembly went back to the baseline, thanks to plastically strengthened I-to-All connections within the assembly.

**Fig 2.**
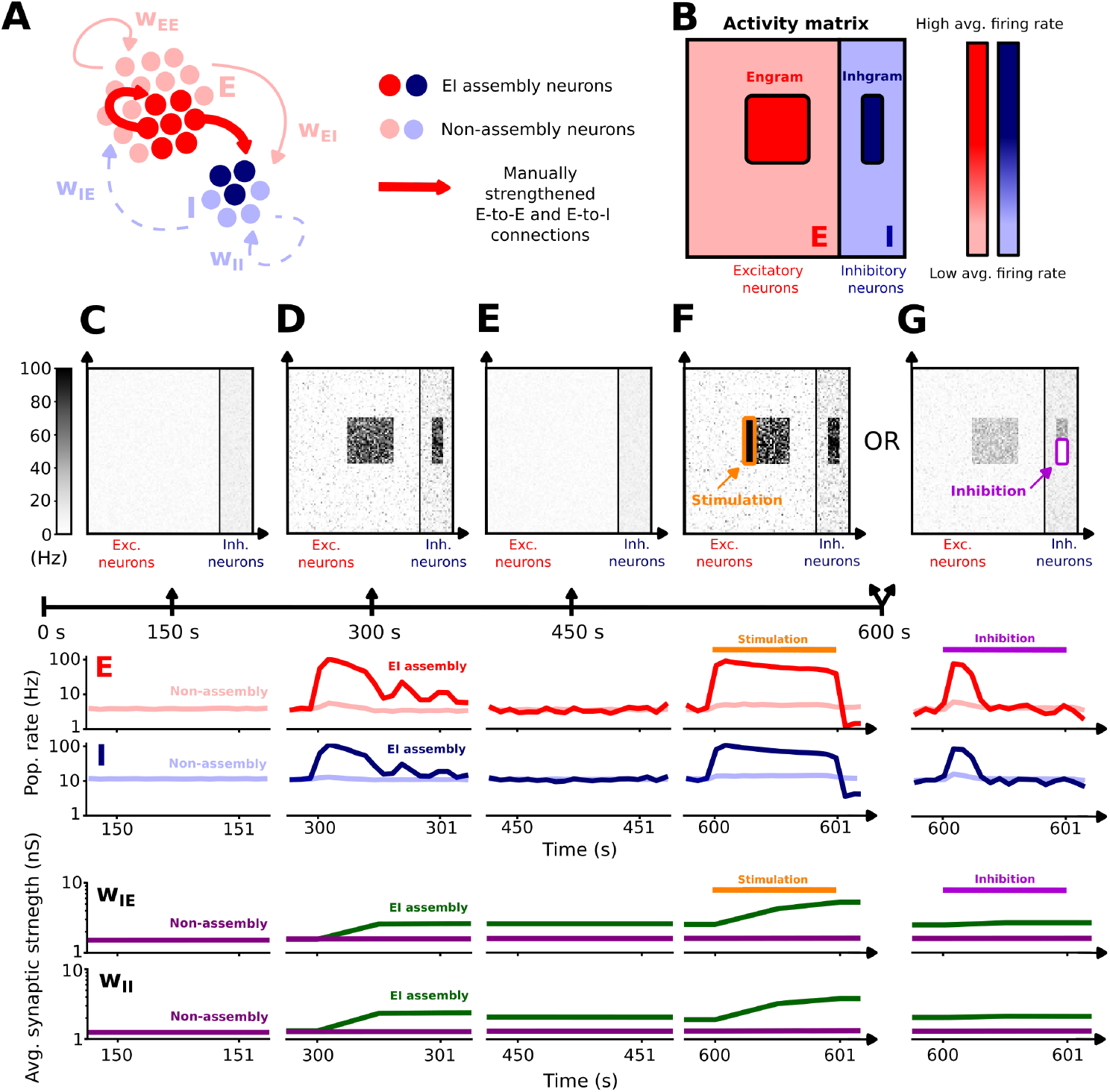
EI assembly self-stabilizes and can be recalled by either stimulation or disinhibition. **A:** Schematic of an RSN with the manually embedded EI assembly. **B:** Schematic of an activity matrix of excitatory and inhibitory populations with visible activity of EI assembly components. Each dot in the matrix represents the activity of a single neuron. **C, D, E, F, G:** Protocol of a recall of the embedded EI assembly. In the stable RSN (C), we manually strengthened the E-to-E and E-to-I connections within a small subpopulation of excitatory and inhibitory neurons (D). After a stabilization process (E), we recalled the EI assembly either by providing partial stimulation to the Engram part (F), or by partially silencing a part of the Inhgram (G). First row: Activity matrices over 0.5 seconds at four different time points (150, 300, 450, 600 seconds). Second row: Population firing rate of excitatory (upper panels) and inhibitory (lower panels) neurons with distinction between the EI assembly sub-population (dark red and dark blue) and non-assembly neurons (in light colours). Third row: Average synaptic strength of I-to-E (*w*_*IE*_ , upper panels) and I-to-I (*w*_*II*_ , lower panels) synapses within the EI assembly (green) and non-assembly (purple) populations.

Finally, we tested engram recall with two protocols. First, we provided a partial cue by stimulating 25% of the engram and monitoring the remaining part of the assembly, as well as the rest of the network (Fig. 2F). The non-stimulated assembly neurons showed elevated activity, indicating successful recall. Second, we tested for the recall by disinhibition. We silenced 50% of the inhgram (inhibitory part of the assembly) and monitored the engram neurons (Fig. 2G). Silencing inhgram neurons successfully increased engram activity for the duration of the modulation, while non-assembly neurons remained largely unaffected.

To further investigate how inhibitory synapses stabilize the EI assembly and enable the recall, we analysed I-to-All connectivity within and outside the EI assembly after embedding (Fig. 3A). We found stronger synaptic weights among EI assembly neurons (I-to-E and I-to-I) compared to non-assembly synapses (Fig. 3B-C). Moreover, the distribution of strengths of synapses within the EI assembly was distinct from the non-assembly connectivity (Fig. 3D). To disentangle the contribution of the I-to-I plasticity, we embedded the same assembly in the network with I-to-E-only^22^ (Fig. 3E). Similar to the I-to-All network, we found increased synaptic strength within the EI assembly connectivity, compared with the nonassembly connections (Fig. 3F-G). However, the distributions of the EI assembly and non-assembly synapses largely overlapped (Fig. 3H), indicating that both populations feature strong weights that could bias the recall of the engram.

**Fig 3.**
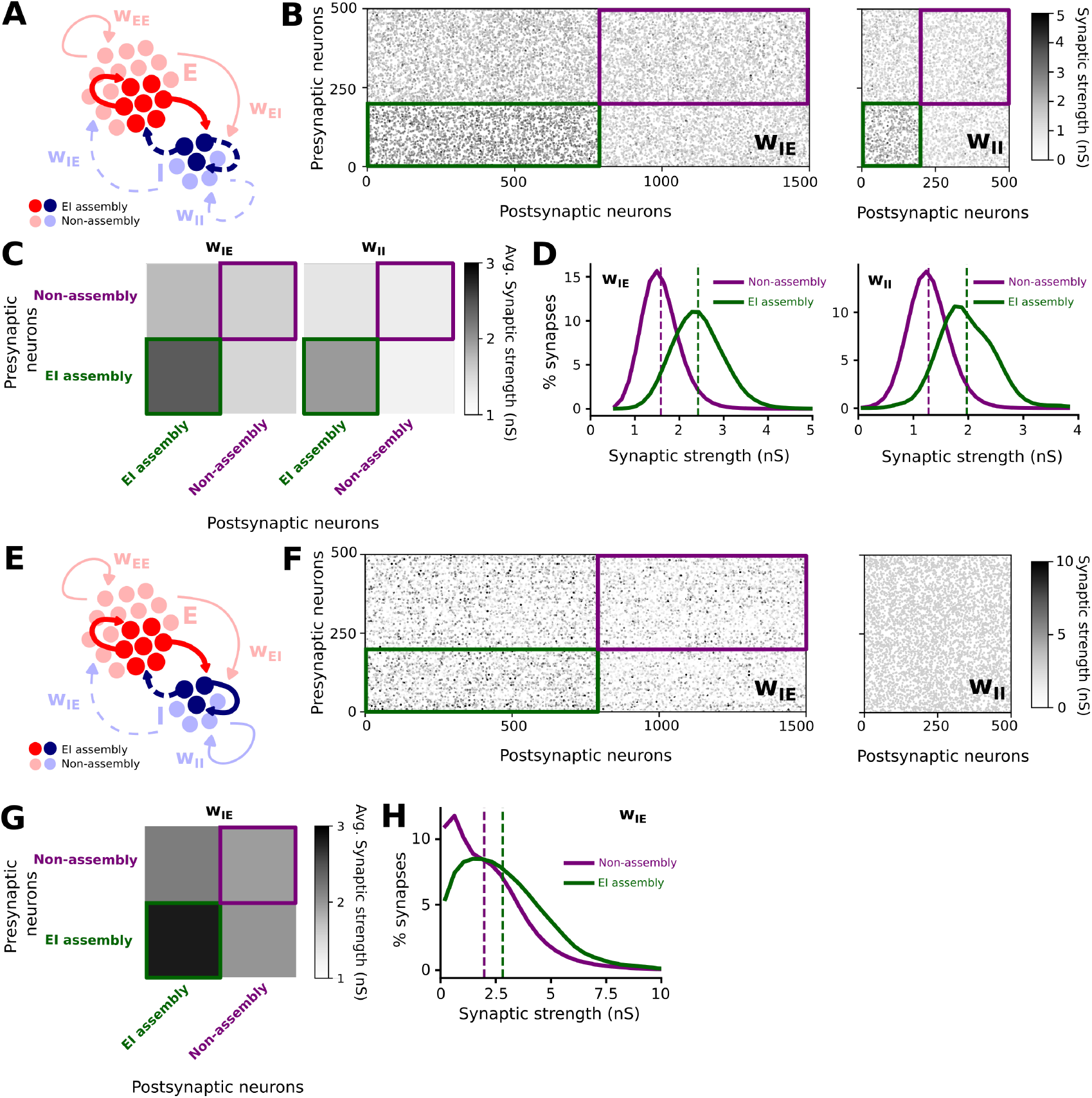
Co-active I-to-I plasticity carves the I-to-E connectivity, making EI assemblies distinguishable. **A:** Schematic of an RSN with the EI assembly and all inhibitory connections plastic. **B:** Examples of connectivity matrices for the I-to-E (left) and I-to-I (right) neurons within the EI assembly (green) and outside of the assembly (purple), with the corresponding synaptic strength. **C:** Average synaptic strength for the I-to-E (*w*_*IE*_ , left) and I-to-I (*w*_*II*_ , right) synapses. **D:** Distribution of *w*_*IE*_ (left) and *w*_*II*_ (right), within the EI assembly (green) and outside of the assembly (purple). **E:** Schematic of an RSN with the EI assembly and only I-to-E connections plastic. **F:** Examples of connectivity matrices for the I-to-E (left) and I-to-I (right) neurons within the EI assembly (green) and outside of the assembly (purple), with the corresponding synaptic strength. Note, I-to-I connections have fixed strength. **G:** Average synaptic strength for the I-to-E (*w*_*IE*_) synapses. **H:** Distribution of *w*_*IE*_ , within the EI assembly (green) and outside of the assembly (purple).

### Pattern completion of an excitatory-inhibitory assembly

Next, we examined its sensitivity to the size of the partial cue in both stimulation-driven (Fig. 4) and disinhibition-driven (Fig. 5) pattern completion protocols. First, we varied the number of stimulated neurons (Fig. 4A-B) and measured population response in networks with I-to-All plasticity (Fig. 4C) and with I-to-E plasticity only (Fig. 4D). We found that both networks could successfully reactivate the EI assembly as indicated by a high signal-to-noise ratio (SNR) (Fig. 4E-F). However, recall in the I-to-All network was more sensitive to stimulation, with an average SNR closer to 0.5 even with small numbers (1%, i.e., 7–8 neurons) of stimulated neurons in the engram. Moreover, SNR was higher in the I-to-All network for all partial-cue sizes.

**Fig 4.**
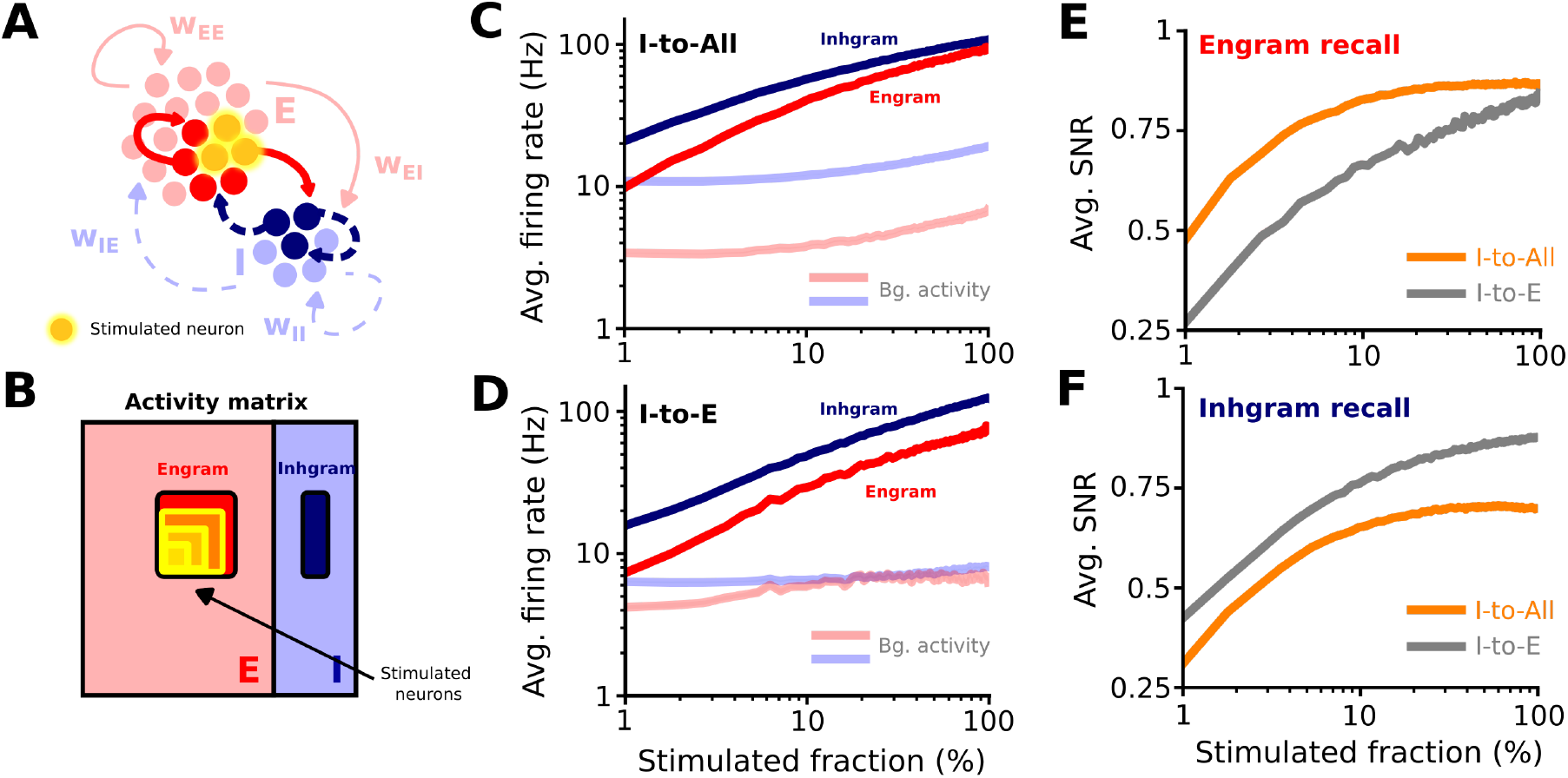
Co-active ISP enhances stimulation-driven engram recall. **A:** Schematic of a direct stimulation of an engram. **B:** Activity matrix representation of the protocol. A partial cue (with increasing size) is provided to the excitatory assembly (engram) to recall the assembly. **C and D:** Average firing rate of Engram (red) and Inhgram (dark blue) populations, as well as background activity (in light colours) for different sizes of the stimulation fraction of the engram, in the I-to-All (C) and I-to-E (D) networks. **E and F:** Average signal-to-noise ratio (SNR) during the recall of the engram (E) and the inhgram (F) for different stimulated fractions of the engram, in the I-to-All (orange) and I-to-E (grey) networks.

**Fig 5.**
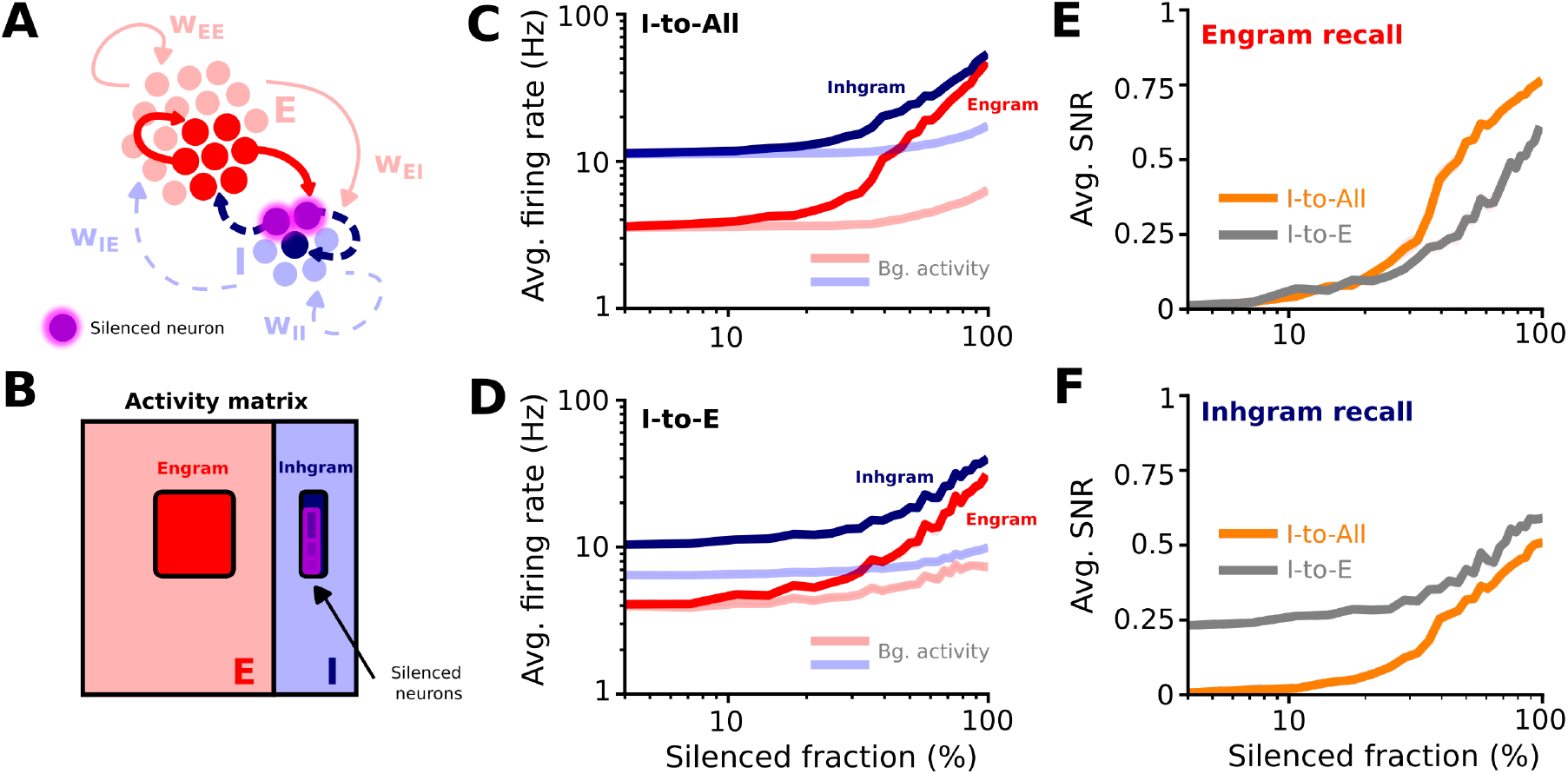
Co-active ISP enhances disinhibition-driven engram recall. **A:** Schematic of silencing an inhgram. **B:** Activity matrix representation of the protocol. A partial inhibition (with increasing size) is provided to the inhibitory assembly (inhgram) to disinhibit the engram. **C and D:** Average firing rate of Engram (red) and Inhgram (dark blue) populations, as well as background activity (in light colours) for different sizes of the silenced fraction of the inhgram, in the I-to-All (C) and I-to-E (D) networks. **E and F:** Average signal-to-noise ratio (SNR) during the recall of the engram (E) and the inhgram (F) for different silenced fractions of the inhgram, in the I-to-All (orange) and I-to-E (grey) networks.

Next, we tested a second recall protocol by varying the number of silenced inhgram neurons (Fig. 5A-B). Increasing the number of silenced neurons elevated assembly activity in both networks (Fig. 5C-D). However, the engram recall was much clearer with silencing fewer neurons in the case of the I-to-All plasticity (Fig. 5E); the average SNR reached approximately 0.5 with 40% (about 78 neurons) of inhibitory engram neurons silenced, whereas in the control network 70% of inhibitory neurons (about 137 neurons) had to be silenced to reach similar SNR levels. Together, our results indicate that a combination of simultaneously active ISP rules on both inhibitory synapse types in the network produced stronger and more sensitive stimulation- and disinhibition-driven recall.

### Recall separation of excitatory-inhibitory assemblies

Single neurons may be a part of multiple engrams^1,24^, so we wondered whether co-active ISP rules could recall overlapping EI assemblies more reliably. First, we tested pattern separation during the stimulation-driven recall (Fig. 6). During this protocol, we overlapped either the engram or inhgram components of two EI assemblies, and stimulated 25% of the first EI assembly. We computed the average SNR between the activities of the two engrams (and, separately, between the two inhgrams) (Fig. 6A-C). Next, we computed the difference between the SNRs (ΔSNR) of the I-to-All and I-to-E networks for engram and inhgram separation (Fig. 6D). We found that both types of networks could successfully separate EI assemblies, with the I-to-All networks performing better when separating engrams with overlap smaller than 50%, and performing equally or slightly worse than I-to-E networks beyond that value.

**Fig 6.**
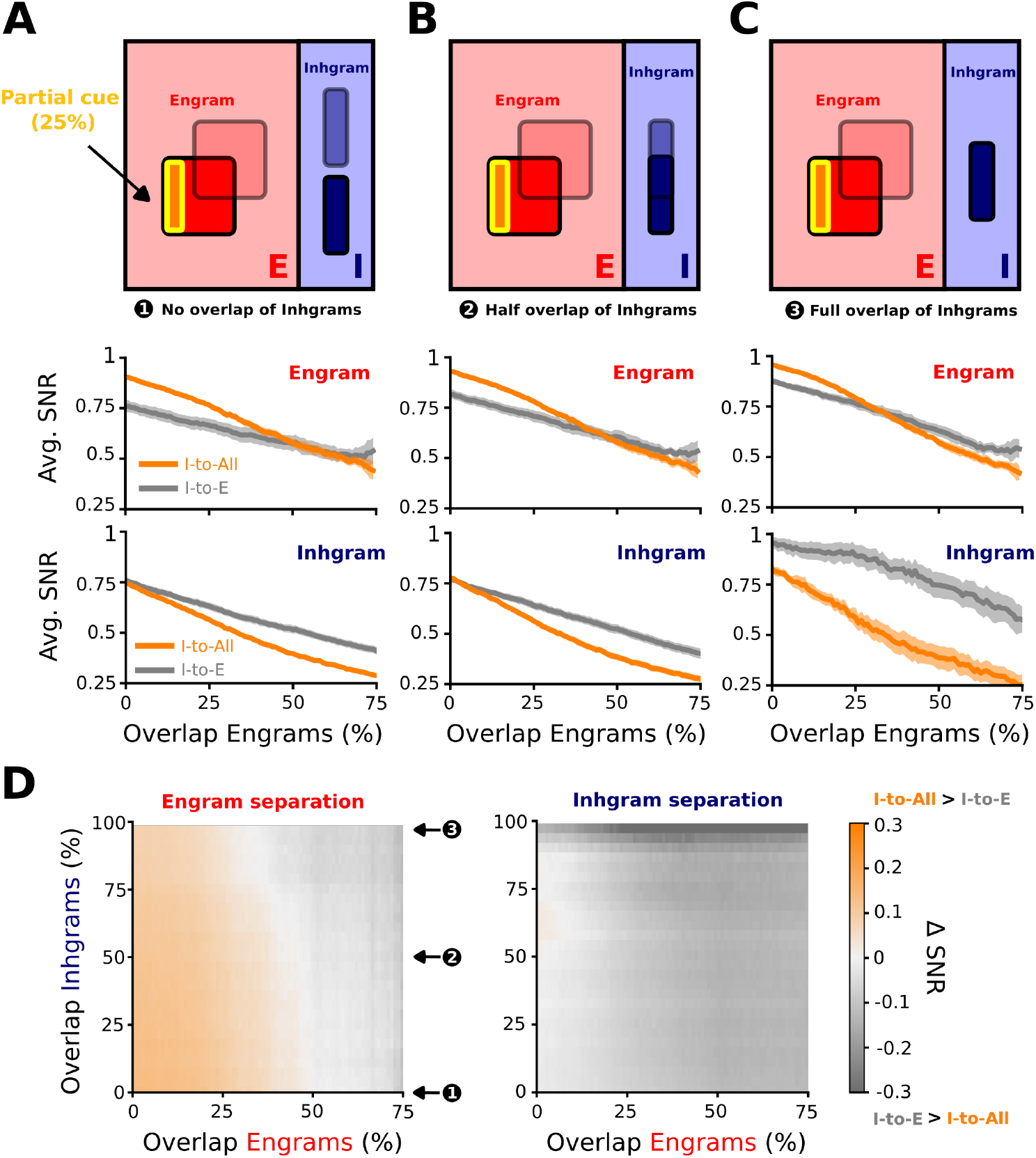
Co-active ISP improves stimulation-driven recall of overlapping engrams. **A, B, and C:** Examples of two EI assemblies with non-overlapping inhgrams (A), half-overlapping inhgrams (B), and fully overlapping inhgrams (C) during the stimulation-driven recall. First row: Activity matrices representing the experiment. Second and third row: Average SNR computed between the activities of two engrams and between the activities of two inhgrams, respectively. **D:** Difference in SNR (ΔSNR) between I-to-All and I-to-E networks during the engram (left) and inhgram (right) separation.

Next, we computed the same metrics but for disinhibition-driven recall (Fig. 7). Here, we also overlapped two EI assemblies; however, we silenced 50% of the inhgram to trigger disinhibition-driven activation of the engram (Fig. 7A-C) and compared ΔSNR between I-to-All and I-to-E networks (Fig. 7D). Importantly, we found that only the I-to-All networks were able to successfully separate both engram and inhgram components, with up to 25% engram overlap. Our results indicate that the simultaneous engagement of ISP rules may facilitate recall of overlapping engrams and facilitate their recall via disinhibitory circuits.

**Fig 7.**
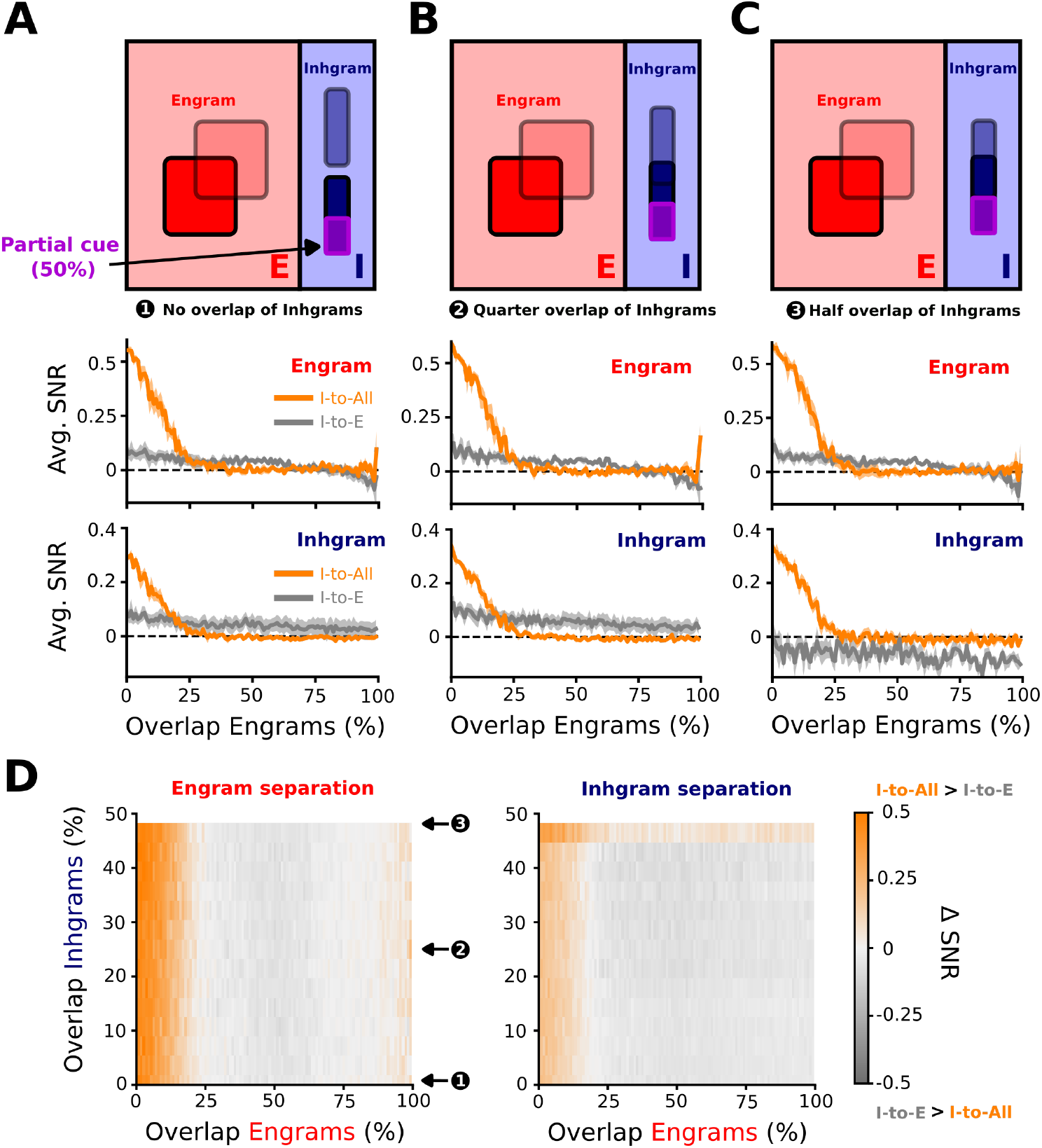
I-to-All plasticity enables disinhibitory recall of overlapping engrams. **A, B, and C:** Examples of two EI assemblies with non-overlapping inhgrams (A), quarter-overlapping inhgrams (B), and half-overlapping inhgrams (C) during the disinhibition-driven recall. First row: Activity matrices representing the experiment. Second and third row: Average SNR computed between the activities of two engrams and between the activities of two inhgrams, respectively. **D:** Difference in SNR (ΔSNR) between I-to-All and I-to-E networks during the engram (left) and inhgram (right) separation.

### Emergence of a functional inhgram in response to an engram embedding

Until this point, we have selected both excitatory and inhibitory members of the EI assembly manually, but we wondered if inhibitory assemblies could emerge autonomously if we chose only the excitatory assembly (Fig. 8A). To detect a potential inhgram, we ranked all inhibitory neurons by their average synaptic weight onto the engram neurons (*w*_I-to-Engram_) and defined candidate inhgrams of increasing size by thresholding this distribution (Fig. 8B). Averaged within each candidate inhgram subpopulation, these connections were stronger than in the non-inhgram population (Fig. 8C). We next asked whether these neurons were also strongly interconnected, by measuring their average outgoing I-to-I weight within each subpopulation (Fig. 8D) and the ratio of outgoing-within to outgoing-outside I-to-I weight (Fig. 8E). We found that neurons that strongly projected to the engram were also strongly interconnected compared with the rest of the population (Fig. 8D-E). We performed the same analysis in the network with I-to-E plasticity only and could identify a corresponding set of neurons with the strongest connections onto the engram (Fig. S5A-C).

**Fig 8.**
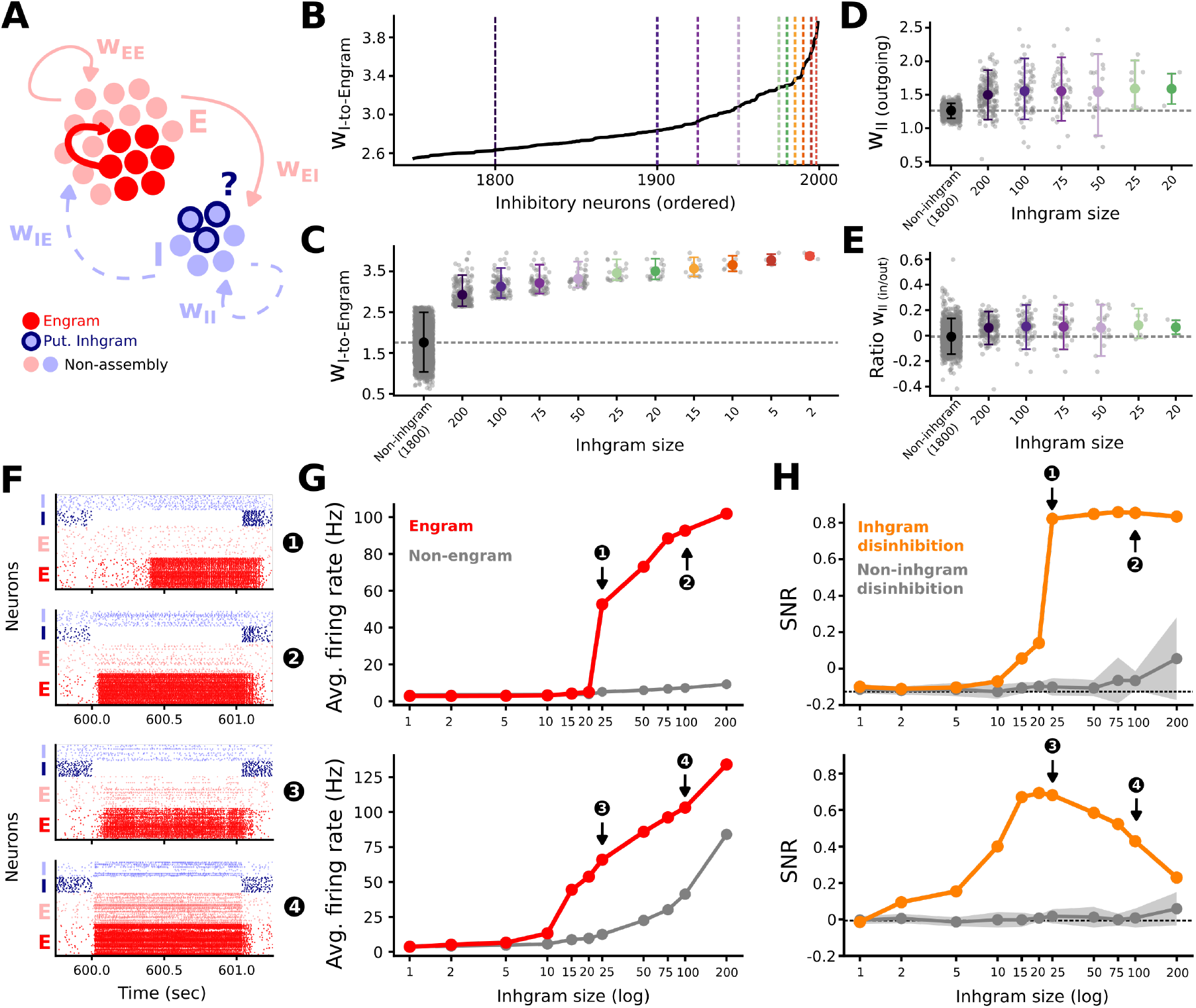
Embedding an engram alone is enough to recruit a functional inhgram. **A:** Schematic of an embedded excitatory engram and the inhibitory neurons potentially recruited as its inhgram. **B:** Average inhibitory-to-engram synaptic weight (*w*_I-to-Engram_) per inhibitory neuron, sorted in ascending order. Vertical dashed lines mark the weight thresholds that define inhgram size. **C:** Average *w*_I-to-Engram_ grouped by inhgram size. The non-inhgram population in grey. **D:** Average I-to-I weight per inhibitory neuron (*w*_*II*_ , outgoing) calculated within each subpopulation. **E:** Ratio of outgoing within and outgoing outside average I-to-I weight (*w*_*II*_ , in/out). **F:** Raster plots of network activity during the disinhibitory recall protocol for the four conditions: two for I-to-All network (1-2), and two for I-to-E network (3-4). **G:** Average firing rate of engram (red) and non-engram (grey) excitatory neurons as a function of inhgram size in both, I-to-All (upper panel) and I-to-E network (lower panel). **H:** Signal-to-noise ratio (SNR) for inhgram disinhibition (orange) and non-inhgram (control) disinhibition (grey; randomly sampled, n=10) as a function of inhgram size in both, I-to-All (upper panel) and I-to-E network (lower panel). Dashed line indicates baseline SNR right before disinhibitory protocol.

Next, we tested whether these emergent inhgrams support disinhibitory recall by silencing inhgrams of increasing size and measuring the SNR during the subsequent engram recall (Fig. 8F-H). Both networks successfully recalled the engram, with the I-to-All network outperforming the I-to-E network, reaching an SNR of approximately 0.86 vs 0.69 (Fig. 8H). This difference was mainly due to stronger activation of the non-engram neurons in the I-to-E network (Fig. 8F). In fact, both networks differed in their sensitivity to the size of the inhibited inhgram. The I-to-All network needed a certain number of neurons to be inhibited to recall an engram, remained stable under large-scale inhibition and retained a high SNR (Fig. 8H). However, the I-to-E network seems to have an optimal region of disinhibitory recall in which enough neurons are inhibited to recall the pattern but not too many so as to destabilize the network’s activity (Fig. 8H). These results suggest that I-to-I plasticity not only drives connectivity that remains stable upon large disruption in global EI balance but also enhances recall of the memory pattern.

Interestingly, the analysis of activity before recall revealed distinct dynamics in the inhibitory populations (Fig. S5D-E). Driven by I-to-E plasticity, excitatory neurons in both networks fired at the target firing rate (3 Hz), with slightly reduced activity of the engram neurons compared to non-engram neurons in the I-to-All network. However, inhgram neurons fired at higher rates than non-inhgram neurons in the I-to-E–only network (Fig. S5E), whereas in the I-to-All network both inhgram and non-inhgram populations, regulated by I-to-I plasticity, fired around the target rate. These results suggest that a fully functional inhgram can be stored in the network while remaining undetectable in the activity of the neurons.

Finally, we asked whether it is possible to predict which neurons will become part of the inhgram before embedding the excitatory assembly. We found that the inhgram neurons already exhibited stronger Engram-to-I (*w*_*EI*_) and I-to-Engram (*w*_*IE*_) connectivity than the rest of the inhibitory population prior to engram embedding in both network types (Fig. 9A-B). This suggests that randomly connected networks may contain pre-existing E–I–E loops that are amplified during the emergence of EI assemblies. We quantified this with a five-fold crossvalidated classifier: the full E–I–E loop (*w*_*EI*_ + *w*_*IE*_) predicted inhgram membership well above chance in both networks, with E-to-I weight (*w*_*EI*_) the dominant predictor in the I-to-All network and I-to-E weight (*w*_*IE*_) dominant in the I-to-E network (Fig. 9C). Together, this result shows that inhgrams involve different circuits during their formation. While I-to-E networks tend to select inhgram neurons through the strong I-to-E connections, the I-to-All networks amplify pre-existing strong E-to-I connections together with plasticity changes on I-to-E and I-to-I connections, forming coupled excitatoryinhibitory assemblies.

**Fig 9.**
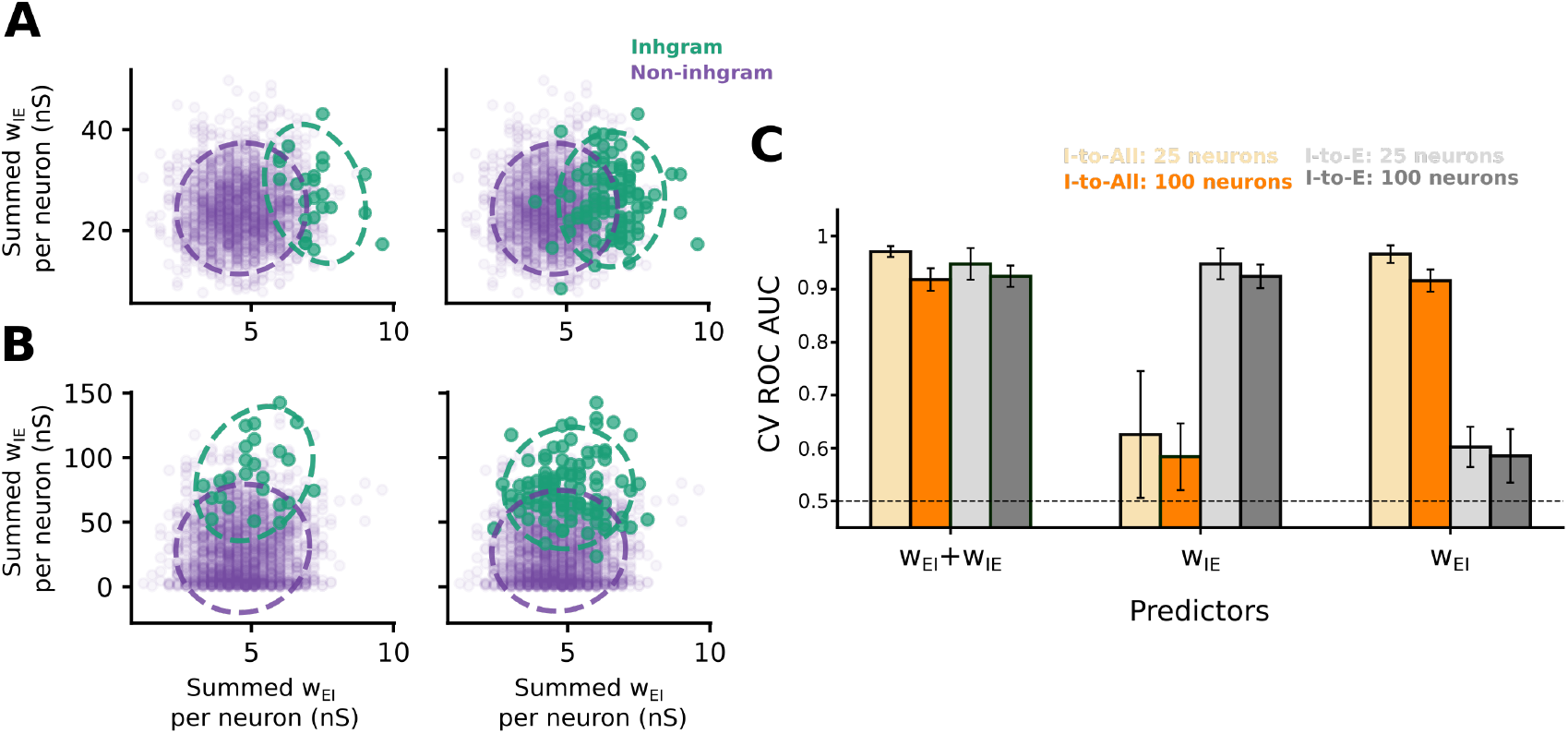
Random connectivity foreshadows which inhibitory neurons become the inhgram. **A:** Joint distribution of the summed I-to-Engram (*w*_*IE*_) and Engram-to-I (*w*_*EI*_) synaptic weight per inhibitory neuron, measured from the random connectivity *before* the engram was embedded, in the I-to-All network. Dashed ellipses summarise the spread of each group. Left: 25-neuron inhgram; right: 100-neuron inhgram. **B:** Same as in A, for the I-to-E network. **C:** Five-fold cross-validated ROC AUC for classifying inhgram membership from the pre-embedding connectivity, using three predictor sets: the full E–I–E loop (*w*_*EI*_ + *w*_*IE*_), the I–E projection alone (*w*_*IE*_), and the E–I projection alone (*w*_*EI*_). Bars show the I-to-All network for 25 (light orange) and 100 (dark orange) inhgram neurons, and the I-to-E network for 25 (light grey) and 100 (dark grey) inhgram neurons; error bars denote standard deviation across folds. Dashed line marks chance.

## III. DISCUSSION

In this work, we show that I-to-I plasticity transforms the role of inhibition from a stabilizing force into an active substrate for memory organisation. Co-active I-to-E and I-to-I plasticity allows inhibitory neurons to not only balance excitatory engrams but to selectively control and reactivate them through disinhibition.

### Shape of the learning rules

We started by analysing the stability of a network with co-active inhibitory synaptic plasticity (Fig. 1), implementing a symmetric rule at both I-to-E and I-to-I synapses. The shape of this rule was introduced theoretically for I-to-E connections^22^ and has some experimental support^23^, but the appropriate form of plasticity for inter-inhibitory connections remains far less constrained. Direct experimental characterization of I-to-I plasticity is still scarce, with only a few exceptions^25–27^. Given its homeostatic effect of driving the firing rates of postsynaptic neurons toward a target rate we considered the rule a good starting point that is also consistent with recent theoretical work showing symmetric shapes as the most prevalent across the space of stability inducing plasticity-rules^28^. A complementary line of work reaches stable excitatory-inhibitory dynamics through cross-homeostatic rules that make both excitatory and inhibitory synapses plastic^29^. In future work, Hebbian and anti-Hebbian inhibitory spike-timingdependent plasticity (STDP) across distinct inhibitory sub-populations, may prove necessary for modular assembly formation^30,31^.

### Contribution of the I-to-I learning to the engram recall

#### Sensitivity to the recall cue

Throughout the study, we tested engram recall under several conditions: the engram as part of an EI assembly, recalled through stimulation (Fig. 4) or disinhibition (Fig. 5), and the engram without an inhibitory component, recalled through stimulation^22^ (Fig. S4). In the EI assembly condition, the I-to-All network performed as well as or better than the I-to-E-only network, and recalled the engram from smaller partial cues. In the (excitatory) engram-only control, without additional inhgram, the two networks performed comparably, indicating that the increased performance of I-to-All plasticity relies on the inhibitory component of the assembly. Moreover, the advantage was specific to recall of the excitatory engram; for the inhibitory component, the I-to-E network produced stronger inhgram recall (Fig. 4F, 5F). Together, these findings suggest that activity-dependent plasticity within the inhibitory population can support more precise and more sensitive recall of stored excitatory memories.

Our findings add to growing work establishing inhibitory dynamics as active contributors to network computation and memory. I-to-E plasticity can shape EI assemblies supporting stimulus-specific computations^17,18^, sculpt complementary inhibitory weight profiles that tune postsynaptic responses^32^, and control the consolidation of neural assemblies through adaptation to selective stimuli^33^, among other functions^10^. Beyond stabilization, recent work shows that disinhibitory circuits can set the sign of excitatory plasticity^34,35^ and that local perturbations of precise EI balance can encode error signals that instruct learning^36^. Finally, co-dependent excitatory–inhibitory plasticity can produce stable, longlasting memory weight profiles^37^. These studies focus on I-to-E plasticity or on disinhibition through fixed inhibitory connectivity. Our results suggest that making I-to-I synapses plastic adds a further layer of organization within the inhibitory population, sharpening and segregating memories, and supporting more precise and quick recall.

#### Selectivity of the engram recall

We also tested selective recall of overlapping EI assemblies (Fig. 6, 7). Under stimulation, the I-to-All network showed only a modest, overlap-dependent advantage (Fig. 6). Under disinhibition, by contrast, co-active plasticity clearly aided the recall of overlapping engrams and outperformed the I-to-E network (Fig. 7) in its discriminating abilities, even when the two assemblies shared half of their inhgram. These findings suggest that co-active inhibitory plasticity enables selective recall specifically through a disinhibitory pathway. Such a feature could be useful in brain networks with a high excitatory-to-inhibitory ratio: the inhibitory population is relatively sparse, around 10–20% of all neurons depending on brain region. We speculate that the excitatory population is suited to storing many distinct memories as engrams, while a comparatively small inhibitory population could retrieve them selectively through disinhibitory circuits that themselves undergo learning, a prediction consistent with the reactivation of human memories by GABA manipulation^13^, making a more fine distinction between discriminating and generalising memory performance.

#### Pre-existing E–I–E loops in the random connectivity

Instead of using *a priori* embedded inhgrams, we also explored if an inhibitory assembly could emerge on its own thanks to plasticity. We could show that such emergent inhgrams formed in both network types, but they were functionally different. Under disinhibitory recall, the I-to-All inhgram produced strong recall as long as the size of silenced inhgram was large enough. The fact that it was invisible at rest suggests that coactive ISP can support memory recall and maintain EI balance at the same time, unlike the I-to-E inhgram (Fig. S5). Interestingly, the inhgram could be predicted from the network’s pre-existing connectivity in both networks, but through different motifs: E-to-I projections dominated the prediction under co-active plasticity and I-to-E projections under I-to-E plasticity (Fig. 9)^38^, indicating that the two rules recruit inhgrams through distinct circuits. Together, these results suggest that coactive ISP sculpts connectivity by amplifying pre-existing E–I–E loops, completing an EI assembly when an engram is encoded. Consistent with this, prior models report that synaptic plasticity can amplify small initial structural biases into organized connectivity^39^, and give rise to distinct assembly types^31^ as well as structured stabilization under I-to-E plasticity^40^.

### Experimental predictions and limitations

#### Heterogeneity of inhibitory population

Our model contains two neuronal populations, one excitatory and one inhibitory, whereas real circuits are far more heterogeneous^41–43^. Our single inhibitory population is best understood as an abstraction of these specialized circuits, but disinhibitory recall has a concrete biological counterpart: SST interneuron-evoked disinhibition in prefrontal cortex can mediate fear memory recall^14^, and dentate gyrus SST interneurons, themselves inhibited by nucleus incertus neurons, mediate fear memory recall as well^15^. It would be informative to understand how real interneuron types implement the inhgrams we describe, and where the model diverges from reality.

We also applied only a single symmetric rule to all inhibitory synapses, but plasticity is unlikely uniform: recent work shows that plasticity rules differ even across the dendritic compartments of individual L2/3 pyramidal neurons in mouse primary motor cortex^44^. Interneuronto-interneuron activity-dependent LTP and LTD remain under-explored, with only a few exceptions^25–27^, yet we expect distinct rules to operate across interneuron classes. A promising way forward is to search for plasticity rules directly in realistic multi-cell-type network models and validate them against in vivo large-scale recordings during a task^45^.

#### Excitatory-inhibitory assemblies in brain networks

We focused on engrams that are coupled to their inhibitory components (inhgrams) through strengthened E-to-I connections (Fig. 2), and we showed that such inhgrams emerge *around* an embedded engram and can be predicted from pre-existing E-I-E loops in both networks (Fig. 8, Fig. 9). The idea that inhgrams might mirror engrams in the brain has been proposed before, particularly in the context of excitatory-inhibitory balance^12^, where balanced EI assemblies could provide targeted feedback inhibition to excitatory neurons. Moreover, such nascent structure doesn’t have to be limited to simple memory assemblies but could also hold true for more elaborate neuronal architectures^46^ that could be sculpted in a “tabula plena”^47^, a rich neural substrate early in development, before underutilised synapses are pruned. Consistently, EI assemblies between excitatory neurons and fast-spiking interneuron subtypes in a zebrafish homologue of piriform cortex may aid associative memory^18^, EI assemblies between pyramidal neurons and PV interneurons may enable stimulus-specific computations in mouse visual cortex^17^, and precisely balanced EI assemblies can shape input representations in sensory cortices^19^, where the spatial structure of inhibition tracks excitatory drive across cortical receptive fields^48^ and control sensory responses in cortical circuits^49^.

Several emerging approaches could test these predictions. Efforts to map connectomes across model organisms, using volumetric electron microscopy in zebrafish^50^ and light-microscopy-based reconstruction combined with deep learning in mammalian cortices^51^, could reveal whether E-I-E loops and disinhibitory motifs are overrepresented in real circuits, as our results predict. In parallel, recording technology continues to advance in precision and scale: Neuropixels electrodes enable simultaneous single-cell-resolution recordings across brain areas over extended periods^52–54^, which could be used to infer functional EI assemblies and track their formation during memory tasks. Finally, precise manipulation of GABA levels in the human brain during fMRI offers a route to test whether inhibition gates memory recall, as our disinhibitory mechanism predicts^13,55,56^.

We propose co-active (or even codependent) inhibitory-inhibitory synaptic plasticity as a mechanistic substrate for disinhibitory memory recall in recurrent spiking networks (RSNs). By engaging plasticity at both I-to-E and I-to-I synapses simultaneously, networks can robustly maintain excitatory-inhibitory assemblies and retrieve them through either direct stimulation or disinhibition. I-to-I plasticity confers a qualitative advantage in the disinhibitory regime, enabling selective pattern separation of overlapping engrams and producing functionally specific inhgrams that can be predicted from pre-existing E-I-E connectivity loops. In summary, these results position inhibitory neurons as active organizers of memory circuits rather than mere stabilizers, and motivate direct experimental tests of disinhibitory recall.

## IV. MATERIALS AND METHODS

### Neuron and network model

We consider a network of 8000 excitatory and 2000 inhibitory neurons, following Vogels, Sprekeler et al.^22^. The membrane potential dynamics of neuron *j* (excitatory or inhibitory) were given by

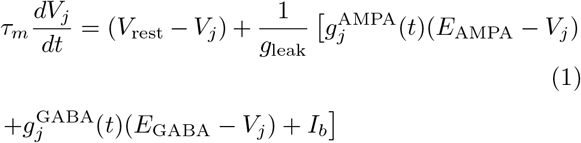

with *τ*_*m*_ = 20 ms, *V*_rest_ = *™*60 mV, *E*_AMPA_ = 0 mV and *E*_GABA_ = *™*80 mV. A postsynaptic spike was emitted whenever the membrane potential *V*_*j*_(*t*) crossed a threshold *V* ^th^ = *™*50 mV, with an instantaneous reset to *V*_rest_ for the duration of the refractory period, *τ*_ref_ = 5 ms. A constant input current *I*_b_ = 200 pA was injected to every neuron to maintain the minimum activity of the network.

The excitatory and inhibitory conductances, 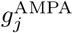 and 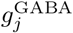, evolved such that:

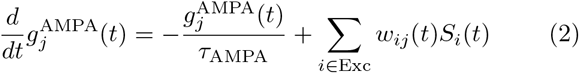

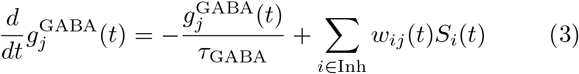

with *τ*_AMPA_ = 5 ms, *τ*_GABA_ = 10 ms, *w*_*ij*_(*t*) the connection strength between neurons *i* and *j* (nS), *S*_*i*_(*t*) =

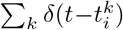 the spike train of presynaptic neuron *i*, where 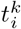 denotes the spike times of neuron *i*, and *δ* the Dirac delta. The recurrent connectivity was instantiated with random sparse connectivity (2%).

### Inhibitory synaptic plasticity (ISP)

Throughout the study, we considered all the excitatory connections fixed with *w*_EE_ = *w*_EI_ = 0.3 nS. The inhibitory-to-excitatory (I-to-E) connections underwent plasticity in both I-to-All and I-to-E networks, while the inhibitory-to-inhibitory (I-to-I) connections underwent plasticity only in I-to-All case but were fixed with *w*_II_ = 3 nS in I-to-E networks. The inhibitory synapses were modified by coincident pre- and postsynaptic activity, following experimentally^23^ and theoretically^22^ described symmetric inhibitory synaptic plasticity rule, with a coincident time window *τ*_STDP_ = 20 ms. The synaptic weight *w*_*ij*_(*t*) from neuron *i* to neuron *j* underwent a Hebbian spike-triggered update for every pre- or postsynaptic event such that

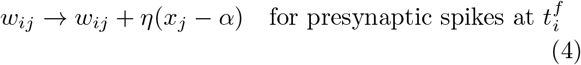

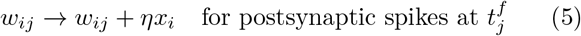

where the learning rate *η* = 1 *×* 10^*™*2^, the depression factor *α* = 2 *× ρ*_0_ *× τ*_STDP_ and *ρ*_0_ is a target firing rate (Hz). Note, we use different *ρ*_0_ for the *w*_IE_ and *w*_II_ connections, which mirrors different target firing rates of excitatory and inhibitory populations.

The synaptic traces *x*_*i*_ and *x*_*j*_ are low-pass filters of the activity of presynaptic neuron *i* and postsynaptic neuron *j*, with time constants *τ*_STDP_, such that

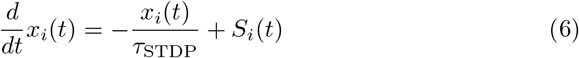

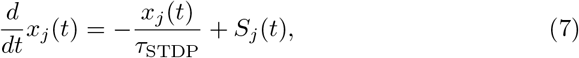

with 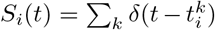 the spike train of neuron *i, δ* the Dirac delta function denoting the presence of a pre-

or postsynaptic spike at time *t*.

### Excitatory-inhibitory assembly recall EI assembly model

In Fig. 2, we introduced a model of an excitatory-inhibitory (EI) assembly. Similarly to Vogels, Sprekeler et al.^22^, the excitatory (engram) and inhibitory (inhgram) assembly were embedded in the network by manually strengthening excitatory connections within a group of 784 excitatory and 196 inhibitory neurons (9.8% of each population) such that 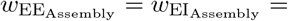 1.5 nS. The EI assembly was embedded after 5 minutes of simulation, once the network had reached a steady state. Recall experiments were performed 5 minutes after embedding the EI assembly (10 minutes after the start of the simulation).

### Pattern completion task

In Fig. 4 and Fig. 5, pattern completion of the embedded EI assembly was tested using two protocols: **stimulation-driven recall** and **disinhibition-driven recall**.

In the stimulation-driven recall condition, the EI assembly was stimulated for 1 second with a partial cue targeting *x*% of the excitatory component of the assembly (engram). The value of *x*% was always chosen as a multiple of 7 neurons, such that the smallest partial cue was 7 neurons (approximately 0.9%) and the largest was 777 neurons (99.1%). The cue was delivered by 1000 Poisson neurons firing at a rate *r*_Poisson_ = 100 Hz, with random sparse connectivity (5%) and synaptic strength *w*_Poisson_ = 3 nS.

Analogously, in the disinhibition-driven recall condition, the EI assembly was inhibited for 1 second using a partial cue targeting *x*% of the inhibitory component of the assembly (inhgram). The value of *x*% was always chosen as a multiple of 7 neurons, such that the smallest partial cue consisted of 7 neurons (approximately 3.6%) and the largest consisted of 189 neurons (96.4%). The cue was delivered by Poisson neurons with the same parameters as in the stimulation-driven condition, except that the input was inhibitory.

The quality of recall in both protocols was assessed using a signal-to-noise ratio (SNR) metric, computed separately for excitatory (SNR_E_) and inhibitory (SNR_I_) part of the assembly, and defined as

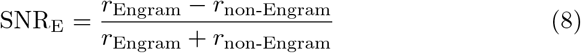

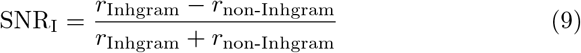

where *r*_Engram (Inhgram)_ denotes the mean activity of neurons that belonged to the engram (inhgram), and *r*_non-Engram (Inhgram)_ denotes the mean activity of neurons outside the assembly. Each condition was simulated ten times with different seeds.

### Pattern separation task

In Fig. 6 and Fig. 7, the pattern separation of two embedded EI assemblies is shown. Here again, it was tested using two protocols: **stimulation-driven recall** and **disinhibition-driven recall**. In both protocols, the EI assemblies were the same size as before and embedded at the same time. The overlap between the engram and inhgram parts of the assemblies was varied independently, so that the EI assemblies could overlap only on engrams, only on inhgrams, or on both.

The cue was delivered the same way as for the pattern completion, and only to the non-overlapping neurons of one of the EI assemblies. In the case of the stimulation-driven recall, 196 (25%) engram neurons were stimulated, and in the case of the disinhibition-driven recall, 98 (50%) inhgram neurons were inhibited.

The quality of separation in both protocols was assessed using a signal-to-noise ratio (SNR) metric, computed separately for excitatory (SNR_E_) and inhibitory (SNR_I_) part of the assembly, and defined as

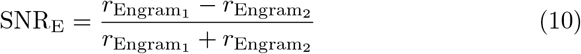

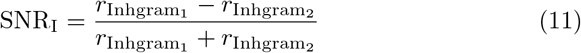

Where 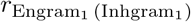 denotes the mean activity of neurons that belonged to the engram (inhgram) of the recalled assembly, and were not directly stimulated (inhibited), and 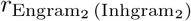 denotes the mean activity of neurons of the non-recalled assembly. Each overlap condition was simulated five times with different seeds.

### Detecting an inhibitory assembly

In Fig. 8A-H, only the engram part (by strengthening only 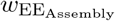) was embedded. 5 minutes after embedding the engram, all inhibitory neurons were ranked by their average synaptic weight onto the engram neurons (*w*_I-to-Engram_), and candidate inhgrams of increasing size were defined by thresholding this distribution (Fig. 8B-C). For each candidate inhgram, we additionally measured the average outgoing I-to-I weight and the ratio of I-to-I weights within to I-to-I weights outside an assembly (Fig. 8D-E). Disinhibitory recall was tested by silencing the candidate inhgram and measuring the resulting engram activity, across the range of inhgram sizes (Fig. 8F- H). As a control, the same number of randomly sampled non-inhgram inhibitory neurons were silenced (*n* = 10 random draws per inhgram size). The SNR_E_ was calculated the same way as for the pattern completion task. The baseline SNR_E_ was calculated over 1 second before the inhibition.

### Predicting inhgram neurons from the pre-existing connectivity

In Fig. 9A-B, the connectivity in the network 5 min after initialization, and right before the engram embedding, was analysed. The synaptic strength of E-to-I and I-to-E were summed per inhibitory neuron and compared against each other for two groups—neurons that will later become an inhgram and the rest of inhibitory population. In Fig. 9C, we quantified how well pre-existing connectivity predicts inhgram neurons using a supervised classification approach. For each inhibitory neuron, three sets of predictors were constructed: (i) the summed E–I– E loop strength, defined as the sum of E-to-I and I-to-E synaptic weights (*w*_*EI*_ +*w*_*IE*_), (ii) the total I-to-E projection strength (*w*_*IE*_), and (iii) the total E-to-I projection strength (*w*_*EI*_). Using these predictors, a binary classifier was trained to distinguish inhgram neurons from the remaining inhibitory population, performed separately for inhgrams of 25 and 100 neurons and for both the I-to-All and I-to-E networks. Prediction performance was evaluated using 5-fold cross-validation, where neurons were randomly split into training and test sets. For each fold, the classifier was trained on 80% of the neurons and evaluated on the remaining 20%. Model performance was quantified using the area under the receiver operating characteristic curve (ROC AUC).

### Data and materials availability

All the code for running the networks and reproducing the analysis is available on GitHub https://github.com/VogelsLab/inhgrams.

## ACKNOWLEDGMENTS

We would like to thank Chaitanya Chintaluri, Douglas Feitosa Tomé, Soumya Kesavabhotla and other Vogels group members for insightful discussions and feedback. We also thank Lisa Topolnik for the support.

## SUPPORTING INFORMATION

**Fig S1.**
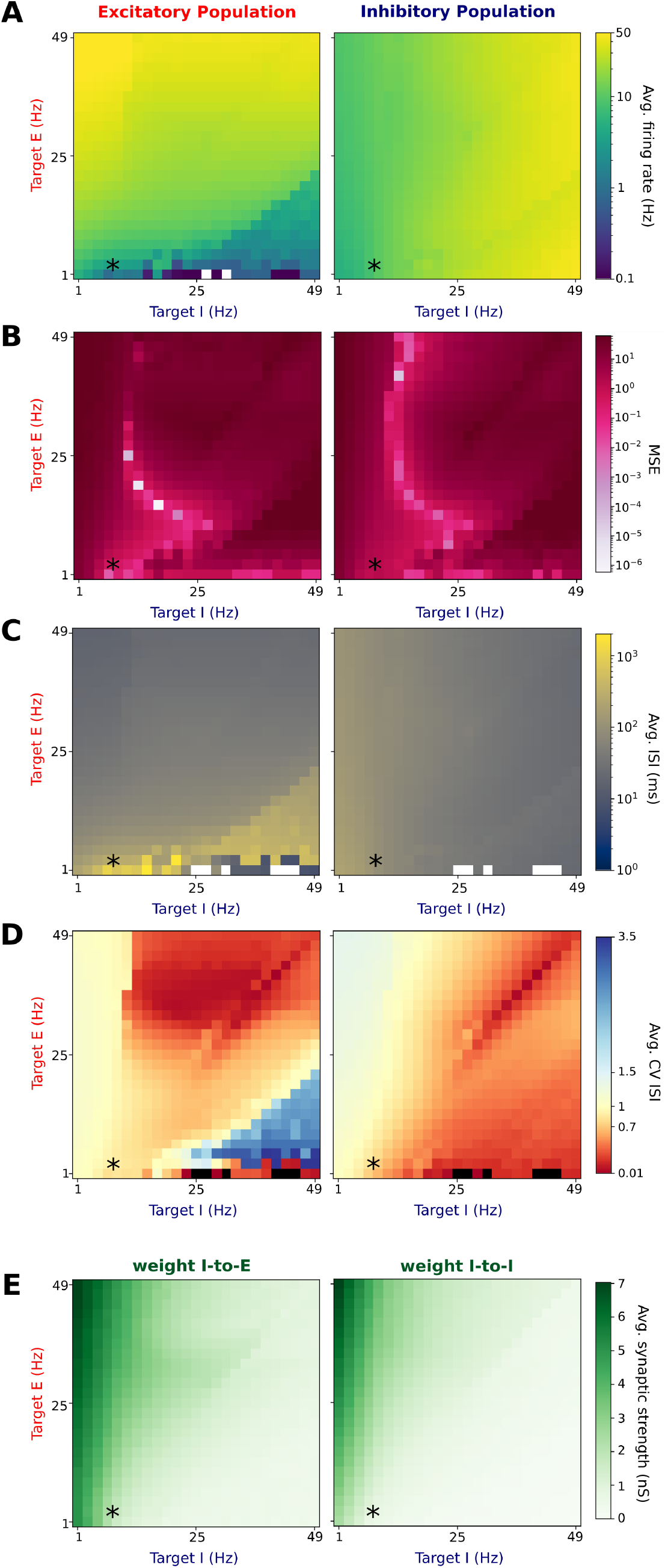
Parameter search over target firing rates of the co-active ISP. All the metrics are computed over the 10 second period after 5 min of network simulation. Left column for excitatory population and right column for inhibitory population. An asterisk shows the network used throughout the paper. **A:** Average firing rate. **B:** Mean squared error (MSE) based on the average firing rate and target firing rate. **C:** Average inter-spike intervals (ISIs). **D:** Average coefficient of variation (CV) of ISIs. **E:** Average synaptic strength of I-to-E (*w*_*IE*_) and I-to-I (*w*_*II*_) connections.

**Fig S2.**
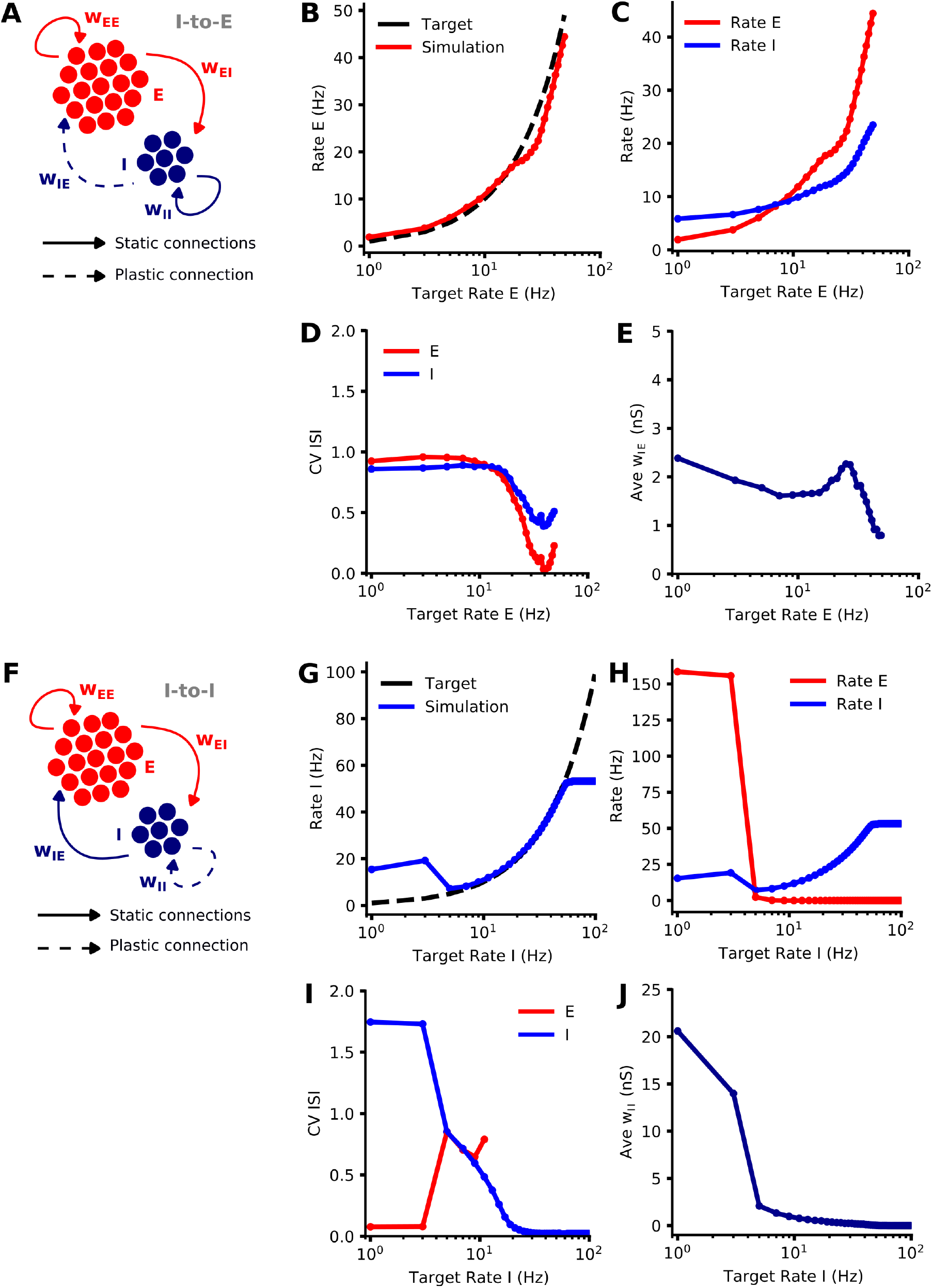
Parameter search over target firing rates of the I-to-E and I-to-I synaptic plasticity, separately. Fixed connections were simulated with following: *w*_*EE*_ = *w*_*EI*_ = 0.3 nS, for A-E: *w*_*II*_ = 3 nS, for F-J: *w*_*IE*_ = 3 nS. **A and F:** Schematics of the network. **B and G:** Average firing rate of excitatory population (panel B) or inhibitory population (panel G) and its target firing rate. **C and H:** Average excitatory and inhibitory rates. **D and I:** Average coefficient of variation (CV) of ISIs. **E and J:** Average synaptic strength of I-to-E (*w*_*IE*_) (panel E) and I-to-I (*w*_*II*_) (panel J) connections.

**Fig S3.**
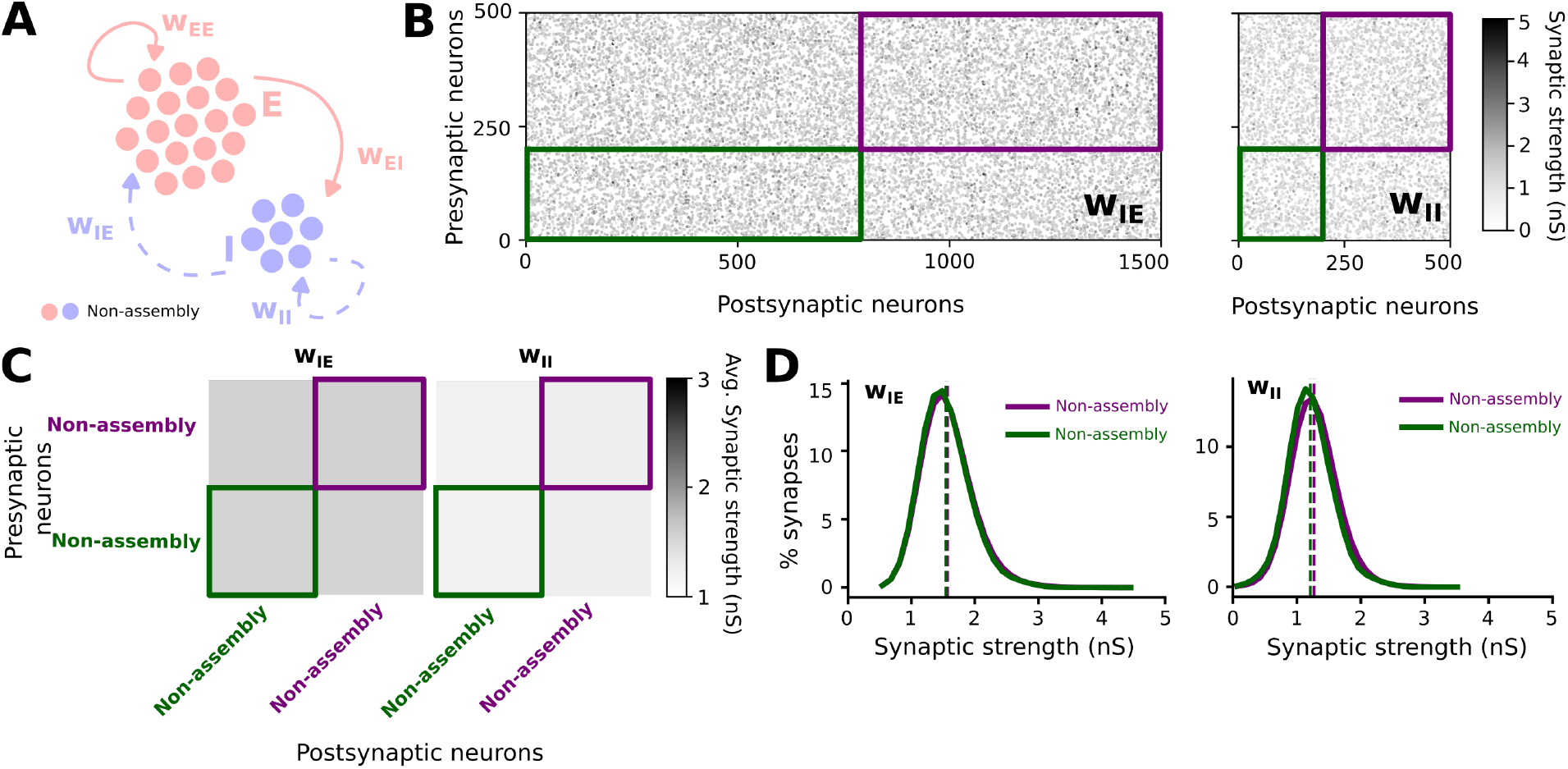
Connectivity of the network with I-to-All plasticity without the excitatory-inhibitory assembly. **A:** Schematic of an RSN without the EI assembly and all inhibitory connections plastic. **B:** Examples of connectivity matrices for the I-to-E (left) and I-to-I (right) neurons within the neurons that would otherwise become an assembly (green) and outside of the assembly (purple), with the corresponding synaptic strength. **C:** Average synaptic strength for the I-to-E (*w*_*IE*_ , left) and I-to-I (*w*_*II*_ , right) synapses. **D:** Distribution of *w*_*IE*_ (left) and *w*_*II*_ (right).

**Fig S4.**
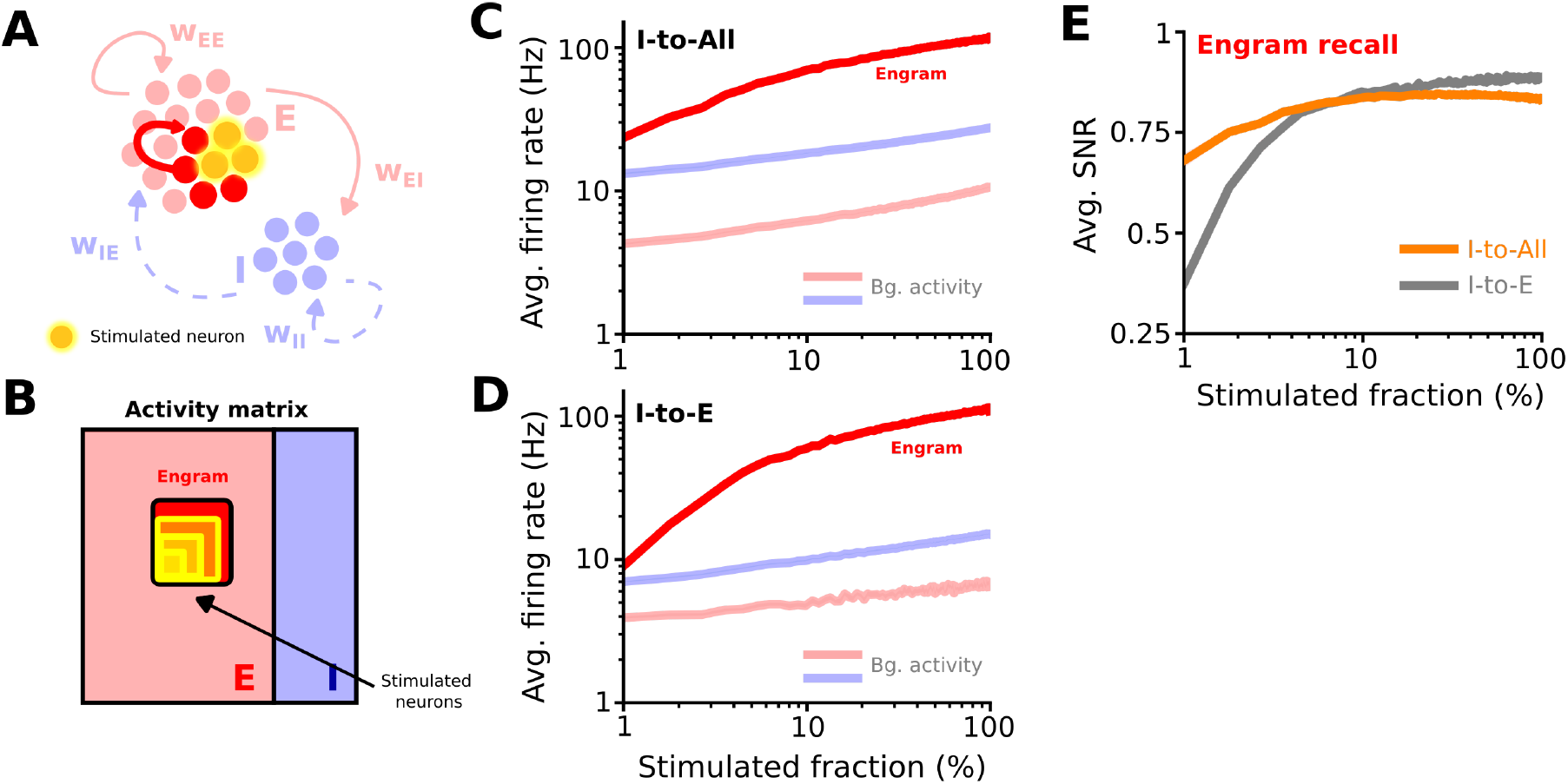
Stimulation-driven engram-only recall in the networks with co-active ISP. **A:** Schematic of a direct stimulation of an engram. **B:** Activity matrix representation of the protocol. A partial cue (with increasing size) is provided to the excitatory assembly (engram) to recall the assembly. **C and D:** Average firing rate of engram (red), as well as background activity (in light colours) for different sizes of the stimulation fraction of the engram, in the I-to-All (C) and I-to-E (D) networks. **E:** Average signal-to-noise ratio (SNR) during the recall of the engram for different stimulated fractions of the engram, in the I-to-All (orange) and I-to-E (grey) networks.

**Fig S5.**
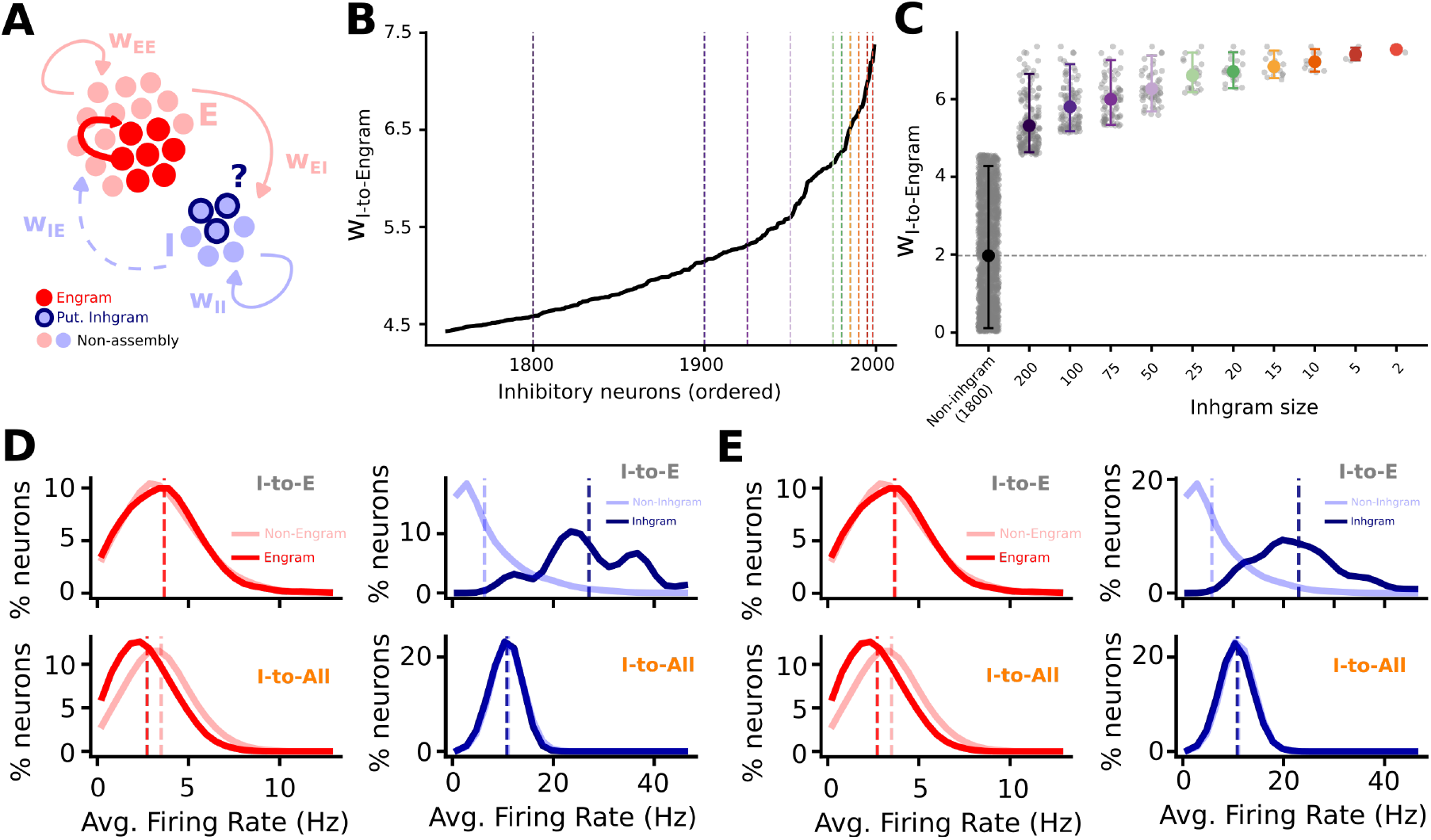
Inhgram detection in the I-to-E plasticity network. **A:** Schematic of a manually embedded engram and potentially recruited inhibitory neurons forming an inhgram. **B:** Average I-to-Engram synaptic weight (*w*_*IE*_) for all inhibitory neurons ordered by connection strength, with dashed vertical lines indicating cutoffs for different inhgram size definitions. **C:** Distribution of *w*_*IE*_ across different inhgram size definitions compared to the non-inhgram population. **D:** Average firing rate distributions of engram (red) and non-engram (light red) excitatory neurons, and inhgram (dark blue) and non-inhgram (light blue) inhibitory neurons (inhgram size 25), in I-to-E (upper panels) and I-to-All (lower panels) networks without perturbations, 5 min after engram embedding. **E:** Same as **D** but for inhgram size 100.

## References

1 S. A. Josselyn and S. Tonegawa, “Memory engrams: Recalling the past and imagining the future,” Science 367, eaaw4325 (2020).

2 C. Van Vreeswijk and H. Sompolinsky, “Chaos in neuronal networks with balanced excitatory and inhibitory activity,” Science 274, 1724–1726 (1996).

3 Y. Shu, A. Hasenstaub, and D. A. McCormick, “Turning on and off recurrent balanced cortical activity,” Nature 423, 288–293 (2003).

4 M. Wehr and A. M. Zador, “Balanced inhibition underlies tuning and sharpens spike timing in auditory cortex,” Nature 426, 442–446 (2003).

5 M. Okun and I. Lampl, “Instantaneous correlation of excitation and inhibition during ongoing and sensory-evoked activities,” Nature neuroscience 11, 535–537 (2008).

6 M. Xue, B. V. Atallah, and M. Scanziani, “Equalizing excitation– inhibition ratios across visual cortical neurons,” Nature 511, 596–600 (2014).

7 M. A. Woodin and A. Maffei, Inhibitory synaptic plasticity (Springer Science & Business Media, 2010).

8 F. Zenke, E. J. Agnes, and W. Gerstner, “Diverse synaptic plasticity mechanisms orchestrated to form and retrieve memories in spiking neural networks,” Nature communications 6, 6922 (2015).

9 G. Hennequin, E. J. Agnes, and T. P. Vogels, “Inhibitory plasticity: balance, control, and codependence,” Annual review of neuroscience 40, 557–579 (2017).

10 Y. K. Wu, C. Miehl, and J. Gjorgjieva, “Regulation of circuit organization and function through inhibitory synaptic plasticity,” Trends in Neurosciences 45, 884–898 (2022).

11 T. P. Vogels and L. Abbott, “Gating multiple signals through detailed balance of excitation and inhibition in spiking networks,” Nature neuroscience 12, 483–491 (2009).

12 H. C. Barron, T. P. Vogels, T. E. Behrens, and M. Ramaswami, “Inhibitory engrams in perception and memory,” Proceedings of the National Academy of Sciences 114, 6666–6674 (2017).

13 H. C. Barron, T. P. Vogels, U. Emir, T. Makin, J. O’shea, S. Clare, S. Jbabdi, R. J. Dolan, and T. Behrens, “Unmasking latent inhibitory connections in human cortex to reveal dormant cortical memories,” Neuron 90, 191–203 (2016).

14 K. A. Cummings and R. L. Clem, “Prefrontal somatostatin interneurons encode fear memory,” Nature neuroscience 23, 61–74 (2020).

15 K. Zichó, K. E. Sos, P. Papp, A. M. Barth, E. Misák, A. Orosz, M. I. Mayer, R. Z. Sebesteny, and G. Nyiri, “Fear memory recall involves hippocampal somatostatin interneurons,” PLoS Biology 21, e3002154 (2023).

16 D. Vallentin, G. Kosche, D. Lipkind, and M. A. Long, “Inhibition protects acquired song segments during vocal learning in zebra finches,” Science 351, 267–271 (2016).

17 O. Mackwood, L. B. Naumann, and H. Sprekeler, “Learning excitatory-inhibitory neuronal assemblies in recurrent networks,” Elife 10, e59715 (2021).

18 C. Meissner-Bernard, B. Jenkins, P. Rupprecht, E. A. Bouldoires, F. Zenke, R. W. Friedrich, and T. Frank, “Computational functions of precisely balanced neuronal microcircuits in an olfactory memory network,” Cell Reports 44 (2025).

19 C. Meissner-Bernard, F. Zenke, and R. W. Friedrich, “Geometry and dynamics of representations in a precisely balanced memory network related to olfactory cortex,” Elife 13, RP96303 (2025).

20 Y. Luz and M. Shamir, “Balancing feed-forward excitation and inhibition via hebbian inhibitory synaptic plasticity,” PLoS computational biology 8, e1002334 (2012).

21 T. P. Vogels, R. C. Froemke, N. Doyon, M. Gilson, J. S. Haas, R. Liu, A. Maffei, P. Miller, C. Wierenga, M. A. Woodin, et al., “Inhibitory synaptic plasticity: spike timing-dependence and putative network function,” Frontiers in neural circuits 7, 119 (2013).

22 T. P. Vogels, H. Sprekeler, F. Zenke, C. Clopath, and W. Gerstner, “Inhibitory plasticity balances excitation and inhibition in sensory pathways and memory networks,” Science 334, 1569–1573 (2011).

23 J. A. D’amour and R. C. Froemke, “Inhibitory and excitatory spike-timing-dependent plasticity in the auditory cortex,” Neuron 86, 514–528 (2015).

24 D. J. Cai, D. Aharoni, T. Shuman, J. Shobe, J. Biane, W. Song, B. Wei, M. Veshkini, M. La-Vu, J. Lou, et al., “A shared neural ensemble links distinct contextual memories encoded close in time,” Nature 534, 115–118 (2016).

25 A. R. McFarlan, C. Guo, I. Gomez, C. Weinerman, T. A. Liang, and P. J. Sjöström, “The spike-timing-dependent plasticity of vip interneurons in motor cortex,” Frontiers in Cellular Neuroscience 18, 1389094 (2024).

26 J. Jab-lońska, G. Wiera, and J. W. Mozrzymas, “Nmda-dependent coplasticity in vip interneuron-driven inhibitory circuits,” Journal of Neurochemistry 169, e70117 (2025).

27 J. Jab-lońska, G. Wiera, and J. W. Mozrzymas, “Non-hebbian long-term depression at vip interneuron inputs selectively tunes inhibition in disinhibitory circuits of mouse hippocampus,” Acta Physiologica 242, e70162 (2026).

28 B. Confavreux, Z. P. Harrington, M. Kania, P. Ramesh, A. N. Krouglova, P. A. Bozelos, J. H. Macke, A. M. Saxe, P. J. Gonçalves, and T. P. Vogels, “Memory by a thousand rules: Automated discovery of multi-type plasticity rules reveals variety & degeneracy at the heart of learning,” bioRxiv , 2025–05 (2025).

29 S. Soldado-Magraner, M. J. Seay, R. Laje, and D. V. Buonomano, “Paradoxical self-sustained dynamics emerge from orchestrated excitatory and inhibitory homeostatic plasticity rules,” Proceedings of the National Academy of Sciences 119, e2200621119 (2022).

30 F. Lagzi, M. C. Bustos, A.-M. Oswald, and B. Doiron, “Assembly formation is stabilized by parvalbumin neurons and accelerated by somatostatin neurons,” BioRxiv , 2021–09 (2021).

31 R. Bergoin, A. Torcini, G. Deco, M. Quoy, and G. Zamora-López, “Emergence and maintenance of modularity in neural networks with hebbian and anti-hebbian inhibitory stdp,” PLoS computational biology 21, e1012973 (2025).

32 E. J. Agnes, A. I. Luppi, and T. P. Vogels, “Complementary inhibitory weight profiles emerge from plasticity and allow flexible switching of receptive fields,” Journal of Neuroscience 40, 9634–9649 (2020).

33 R. Bergoin, A. Torcini, G. Deco, M. Quoy, and G. Zamora-López, “Inhibitory neurons control the consolidation of neural assemblies via adaptation to selective stimuli,” Scientific Reports 13, 6949 (2023).

34 J. Rossbroich and F. Zenke, “Dis-inhibitory neuronal circuits can control the sign of synaptic plasticity,” Advances in Neural Information Processing Systems 36, 64059–64082 (2023).

35 M. Canto-Bustos, F. K. Friason, C. Bassi, and A.-M. M. Oswald, “Disinhibitory circuitry gates associative synaptic plasticity in olfactory cortex,” Journal of Neuroscience 42, 2942–2950 (2022).

36 J. Rossbroich and F. Zenke, “Breaking balance: Encoding local error signals in perturbations of excitation-inhibition balance,” bioRxiv , 2025–05 (2025).

37 E. J. Agnes and T. P. Vogels, “Co-dependent excitatory and inhibitory plasticity accounts for quick, stable and long-lasting memories in biological networks,” Nature Neuroscience 27, 964–974 (2024).

38 E. Giannakakis, O. Vinogradov, V. Buendía, and A. Levina, “Structural influences on synaptic plasticity: The role of presynaptic connectivity in the emergence of e/i co-tuning,” PLOS Computational Biology 20, e1012510 (2024).

39 F. Effenberger, J. Jost, and A. Levina, “Self-organization in balanced state networks by stdp and homeostatic plasticity,” PLoS computational biology 11, e1004420 (2015).

40 D. Festa, C. Cusseddu, and J. Gjorgjieva, “Structured stabilization in recurrent neural circuits through inhibitory synaptic plasticity,” bioRxiv , 2024–10 (2024).

41 H. Markram, M. Toledo-Rodriguez, Y. Wang, A. Gupta, G. Silberberg, and C. Wu, “Interneurons of the neocortical inhibitory system,” Nature reviews neuroscience 5, 793–807 (2004).

42 L. Topolnik and S. Tamboli, “The role of inhibitory circuits in hippocampal memory processing,” Nature Reviews Neuroscience 23, 476–492 (2022).

43 A. Tzilivaki, J. J. Tukker, N. Maier, P. Poirazi, R. P. Sammons, and D. Schmitz, “Hippocampal gabaergic interneurons and memory,” Neuron 111, 3154–3175 (2023).

44 W. J. Wright, N. G. Hedrick, and T. Komiyama, “Distinct synaptic plasticity rules operate across dendritic compartments in vivo during learning,” Science 388, 322–328 (2025).

45 Z. Ye, B. Huang, Y. Wu, G. Chen, and J. Wu, “Discovering heterogeneous synaptic plasticity rules via large-scale neural evolution,” in The Fourteenth International Conference on Learning Representations (2026).

46 T. P. Vogels and L. F. Abbott, “Signal propagation and logic gating in networks of integrate-and-fire neurons,” Journal of neuroscience 25, 10786–10795 (2005).

47 V. Vargas-Barroso, J. F. Watson, A. Navas-Olive, A. Schlögl, and P. Jonas, “Developmental emergence of sparse and structured synaptic connectivity in the hippocampal ca3 memory circuit,” Nature Communications (2026).

48 C. I. Moore and S. B. Nelson, “Spatio-temporal subthreshold receptive fields in the vibrissa representation of rat primary somatosensory cortex,” Journal of neurophysiology 80, 2882–2892 (1998).

49 I. Volkov and A. Galazyuk, “Peculiarities of inhibition in cat auditory cortex neurons evoked by tonal stimuli of various durations,” Experimental brain research 91, 115–120 (1992).

50 R. W. Friedrich and A. A. Wanner, “Dense circuit reconstruction to understand neuronal computation: focus on zebrafish,” Annual Review of Neuroscience 44, 275–293 (2021).

51 M. R. Tavakoli, J. Lyudchik, M. Januszewski, V. Vistunou, N. Agudelo Dueñas, J. Vorlaufer, C. Sommer, C. Kreuzinger, B. Oliveira, A. Cenameri, et al., “Light-microscopy-based connectomic reconstruction of mammalian brain tissue,” Nature 642, 398–410 (2025).

52 V. P. Pastore, P. Massobrio, A. Godjoski, and S. Martinoia, “Identification of excitatory-inhibitory links and network topology in large-scale neuronal assemblies from multi-electrode recordings,” PLoS computational biology 14, e1006381 (2018).

53 N. A. Steinmetz, C. Aydin, A. Lebedeva, M. Okun, M. Pachitariu, M. Bauza, M. Beau, J. Bhagat, C. Böhm, M. Broux, et al., “Neuropixels 2.0: A miniaturized high-density probe for stable, long-term brain recordings,” Science 372, eabf4588 (2021).

54 E. M. Trautmann, J. K. Hesse, G. M. Stine, R. Xia, S. Zhu, D. J. O’Shea, B. Karsh, J. Colonell, F. F. Lanfranchi, S. Vyas, et al., “Large-scale high-density brain-wide neural recording in nonhuman primates,” Nature Neuroscience 28, 1562–1575 (2025).

55 R. S. Koolschijn, U. E. Emir, A. C. Pantelides, H. Nili, T. E. Behrens, and H. C. Barron, “The hippocampus and neocortical inhibitory engrams protect against memory interference,” Neuron 101, 528–541 (2019).

56 R. S. Koolschijn, A. Shpektor, W. T. Clarke, I. B. Ip, D. Dupret, U. E. Emir, and H. C. Barron, “Memory recall involves a transient break in excitatory-inhibitory balance,” Elife 10, e70071 (2021).

